# Evaluating the Link Between Efflux Pump Expression and Motility Phenotypes in *Pseudomonas aeruginosa* Treated with Virulence Inhibitors

**DOI:** 10.1101/2025.01.07.631739

**Authors:** Hannah K. Lembke, Kelsie. M. Nauta, Ryan C. Hunter, Erin E. Carlson

## Abstract

Antibiotic resistance continues to rise as a global health threat. Novel anti-virulence strategies diminish the drive for evolutionary pressure, but still hinder a pathogen’s ability to infect a host. Treatment of the highly virulent *Pseudomonas aeruginosa* strain PA14 with virulence inhibitors (R-2 and R-6) elicited widely varying transcriptional profiles. Of interest, expression of a family of resistance-nodulation-division (RND) efflux pumps implicated in the intrinsic drug resistance of *P. aeruginosa*, was significantly altered by R-2 and R-6 treatment. While structurally similar, these inhibitors caused differential expression of various RND efflux pumps within the Mex family—R-2 treatment stimulated expression of *mexEF-oprN* while R-6 treatment led to increased *mexAB-oprM* expression. Further expansion into a small library of virulence inhibitors revealed chemical motifs that trigger increases in RND efflux pump expression. Additionally, activation of these efflux pumps suggests low accumulation of virulence inhibitors in WT PA14. Treatment of an efflux pump-deficient strain with R-2 or R-6 resulted in inhibition of several virulence factors, for example R-2 was found to abolish swimming motility. Collectively, treatment with either R-2 or R-6 gives rise to a convoluted transcriptomic response, confounded by the impact of efflux pump expression on the system. However, understanding the moieties that lead to high expression of the efflux pumps enables further rational design of novel virulence inhibitors that do not cause RND efflux pump activation.

**Summary:** Antibiotic resistance continues to rise across the world. The continued failure of antibiotics to treat infectious disease clearly demonstrates the need for new therapeutics. Small molecule inhibitors of virulence, or the ability of a bacteria to establish and maintain an infection, have showed promise in this area. By reducing virulence, the host immune system can clear the infection on its own and there is less selective pressure on the bacteria, reducing the emergence of resistance.

Our work demonstrates a clear link between virulence inhibitors and efflux pump expression. Efflux pumps are critical for bacteria to extrude toxic compounds from their interior. Interestingly, the changes in efflux pump expression can be quite large even with similar molecular structures. Additionally, the repression of efflux expression caused by our virulence inhibitors results in synergy with current antibiotics. This work will allow the future design of virulence inhibitors that do not activate efflux pump expression and therefore, enable high intracellular concentrations.

## Introduction

Existing antibiotics inhibit essential cell functions, such as protein or cell wall synthesis, membrane integrity, DNA replication, and metabolic pathways. However, inhibiting essential cell functions leads to high selective pressure for bacteria to evolve antibiotic resistance (ABR). Targeting pathogen virulence has gained popularity as a means to address the increasing rate of failure of many antibiotics and the threat of multi- and pan-drug resistance.(1) By decreasing virulence, an organism’s ability to infect or drive pathogenesis is reduced and is subsequently cleared by the host immune system.

*P. aeruginosa* is a Gram-negative opportunistic pathogen and is difficult to treat due to several compounding features.(2–5) First, the cell envelope of Gram-negative organisms is difficult to permeate with antibiotics due to the presence of a hydrophilic outer membrane (OM) complemented by a hydrophobic inner membrane (IM).(6) Additionally, of the ∼40 porins encoded in the *P. aeruginosa* chromosome, none are non-specific, making compound entry a further challenge.(7, 8) Lastly, *P. aeruginosa* harbors a large family of Resistance-Nodulation-Division (RND) efflux pumps, known for extruding antibiotics resulting in high intrinsic levels of ABR. RND efflux pumps are tripartite pumps comprised of an IM subunit, a periplasmic spanning subunit, and an OM protein that enable them to remove chemicals from either the cytosol or periplasm.(9) The most well-known of the RND efflux pumps is composed of MexAB (IM and periplasmic spanning subunits) and OprM (OM protein). MexAB-OprM is constitutively active in *P. aeruginosa* and extrudes multiple classes of structurally diverse antibiotics and toxic compounds.(10–13) In addition to MexAB-OprM, MexCD-OprJ, MexEF-OprN and MexXY-OprM are largely responsible for the intrinsic ABR of *P. aeruginosa*.(9) Taken together, the impermeable nature of the cell envelope combined with RND-mediated drug efflux, makes intracellular compound accumulation difficult to achieve.

*P. aeruginosa* also wields an impressive virulence arsenal. Virulence mechanisms include toxin production, quorum sensing, adhesion, and invasion (aided by the type II and III secretion systems respectively). Additionally, *P. aeruginosa* tightly regulates its motility, because swimming and swarming enable pathogenesis during acute infections, whereas twitching followed by biofilm formation promotes chronic infections.(2, 14–17) Many virulence phenotypes are regulated by signal transduction systems called two-component systems (TCSs), typically composed of a sensor histidine kinase (HK) and cognate response regulator.(18, 19) Of the 64 annotated HKs in *P. aeruginosa*, 63 are linked to virulence though most are non-essential, making them viable candidates for novel anti-virulence therapies focusing on minimizing ABR.(20) Additionally, HKs have highly conserved catalytic and ATP-binding (CA) domains, enabling pan-inhibition of HKs and potential repression of multiple virulence factors with a single inhibitor.(21) We have previously identified a promising 2-aminobenzothiazole scaffold that targets the CA domain for HK inhibition.(22) Two compounds containing this core, R-2 and R-6, exhibited high *in vitro* potency against a representative HK (IC_50_ = 1.21 µM and 15.1 µM, respectively).(23) Subsequently, both R-2 and R-6 were found to inhibit virulence mechanisms in a pathogenic strain of *P. aeruginosa* (PA14).(24) R-2 is dimer-like in structure with an ether linkage at the 6-position of the ring while R-6 has a -CF_3_ group at this position (**Fig 1A**). Even with high structural similarity, these inhibitors yield different cellular phenotypes with toxin production reduced under R-6 treatment, and swarming motility repressed under R-2 treatment, suggesting R-2 and R-6 may target different HKs.(24)

**Fig 1:**
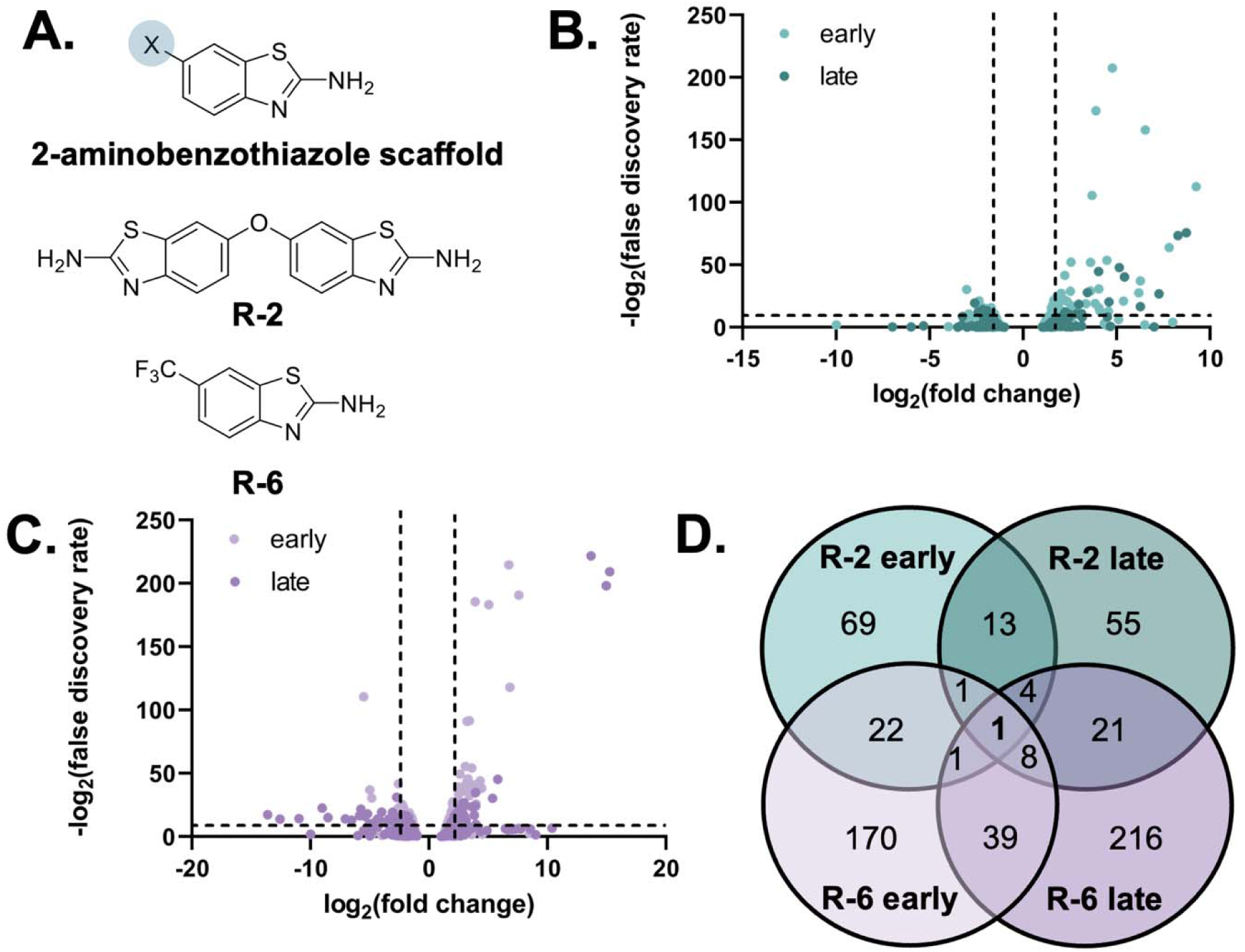
Overview of RNA-Sequencing Results. A. Structure of top HK inhibitor 2-aminobenzothiazole scaffold and inhibitors (R-2 and R-6) used in the RNA-sequencing experiment. B. and C. Volcano plots of expression fold changes compared to DMSO control for R-2 and R-6 treatment, respectively. Dashed bars indicate log_2_(fold change) > 2 on the x-axis and *p*-values < 0.05 on the y-axis. Any genes that met both criteria were considered significantly differentially expressed. D. Venn diagram depicting differentially expressed genes in each data set. Numbers in the overlapped circles indicate genes shared between datasets.

Given these contrasting phenotypes, we hypothesized that treatment with R-2 and R-6 would drive significant transcriptional perturbation. To test this, PA14 was treated with each compound at mid-exponential (OD_600_ ∼0.9, ‘early’) and early stationary phase (OD_600_ 1.2, ‘late’) that corresponded with time points chosen in our previous assays of PA14 virulence.(24) The resulting gene expression profiles revealed unique pathways that were altered by R-2 or R-6 treatment, defined the target variation between these inhibitors, and informed how growth phase influences susceptibility to these compounds. Through this analysis, we found a complicated connection between inhibitor treatment, RND efflux pump expression, and virulence. Informed by our gene expression profiles, we generated efflux pump-deficient strains to evaluate how different efflux pumps impact virulence phenotypes and began to identify structural motifs in HK inhibitors that avoid increasing efflux pump expression.

## Results & Discussion

### RNA-Sequencing Analysis

To define how R-2 and R-6 alter gene expression, PA14 was treated with a DMSO control, R-2, or R-6, at both early (OD600 ∼0.9) and late (OD600 ∼1.2) timepoints, mRNA was extracted and sequenced using the Illumina HiSeq platform. Raw sequence data were analyzed in edgeR (Bioconductor package, R) to identify differentially expressed genes (gene with a fold-change >2 and p-value <0.05) for each condition compared to its DMSO control. Differentially expressed genes from each condition were prioritized for further investigation (**S1 Table**). We found R-2 at the early timepoint (R-2E) altered the expression of 69 genes (49 up, 20 down), whereas at the R-2 late (R-2L) timepoint, expression of 55 genes changed significantly (32 up, 23 down, **Fig 1B**) with 13 genes overlapping between R-2 timepoints (**Fig 1D**). R-6 caused differential expression of 170 and 216 genes at R-6E and R-6L, respectively (**Fig 1C**), with 39 shared. Comparing within early and late timepoints, 22 and 21 genes were shared between the R-2 and R-6 treatments, respectively (**Fig 1D**). A single gene was found in common across all four datasets, *PA14_71100,* annotated as a hypothetical membrane protein with no orthologs in the *Pseudomonas* Genome Database.(25)

We then focused on differentially expressed genes in each condition. Of the 69 transcripts in the R-2E condition, STRING analysis demonstrated that the largest group of related genes were associated with anaerobic growth (**S1 Fig**). Specifically, major genes relevant for the denitrification pathway (*norB, norC, nirS, nosZ*) in PA14 were all upregulated. Interestingly, *narG* and *narK2,* encoding a subunit of the nitrate reductase complex and a nitrite reductase, respectively, were downregulated. Linked to these genes is *coxB,* a cytochrome c oxidase subunit, and *xenB,* a xenobiotic reductase, responsible for reducing nitro-containing compounds. Interestingly, linked to *xenB* in the STRING analysis are the *mexF, mexE*, and other genes found in the *mexT* operon (*PA14_28410, PA14_56640* and *PA14_22420*), but not *mexT* itself. MexT is the native repressor of MexEF-OprN and is redox-responsive.(26) The MexEF-OprN efflux pump is also known to be activated during nitrosative stress in conjunction with XenB.(27)

In the R-2L condition, transcripts related to anaerobic metabolism were not significantly upregulated compared to the DMSO control (**S1 Table, S2 Fig**), but expression of *mexF* and *oprN* remained high (4.3- and 7.0-fold increase, respectively). Additionally, *mexAB-oprM,* encoding the MexAB-OprM efflux pump, and its native repressor *mexR* (**Fig 2A**) were significantly downregulated.(28) Chemotaxis related genes such as *cheW, cheD, cheR2, aer2,* and *PA14_02230* were also repressed while genes related to lipopolysaccharide production (*arnE, arnT* and *PA14_18300*) were upregulated relative to DMSO. Finally, *opmE,* another OM efflux protein within the Mex family (canonically paired with MexPQ) (29) was temporally upregulated.

**Fig 2:**
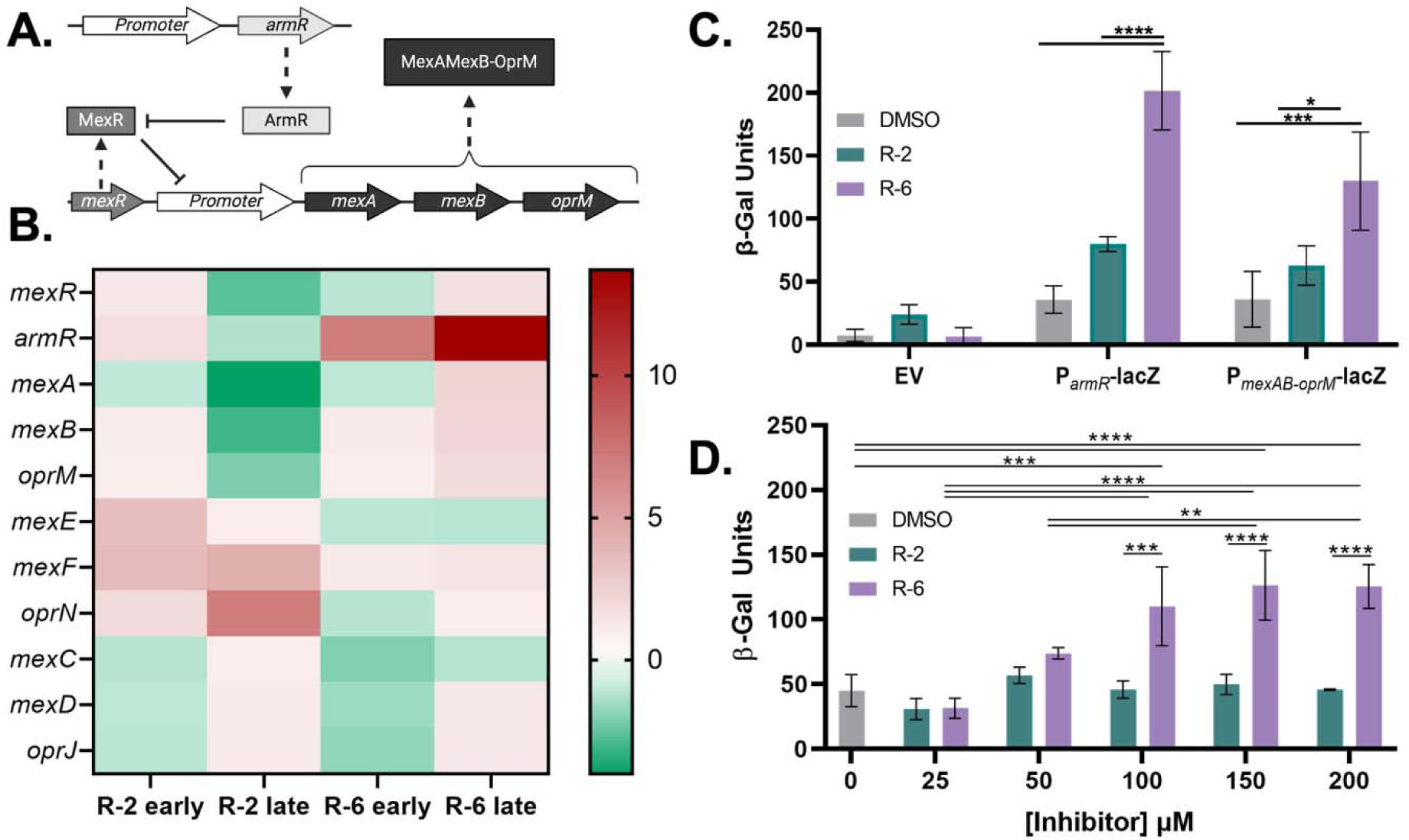
HK Inhibitors Impact on *armR* and *mexAB-oprM* Expression. A. The *mexAB-oprM* operon in PA14. B. Heat map for Mex family efflux pump genes under each treatment condition. C. Validation of RNA-sequencing data was achieved through the generation of P*_armR_-*lacZ and P*_mexAB-oprM_*-lacZ reporter fusions in PA14. R-2 and R-6 were dosed at 200 µM and compared to both a DMSO and empty vector (EV) control. All experiments were completed in technical and biological triplicates. D. Various doses of R-2 or R-6 with the P*_armR_*-lacZ PA14 strain. Statistical significance for panels C and D were determined through a one-way ANOVA. *p*-value * <0.01, ** <0.005, ***<0.0005, ****<0.0001.

Evaluating R-6E compared to DMSO, showcases many interconnecting clusters via STRING analysis (**S3 Fig**). The *pel* genes (*pelA, pelD, pelE, pelF, pelG)* were all upregulated indicating formation of the pel polysaccharide required for PA14 biofilm formation.(30) Accordingly, a large group of *tad* genes responsible for assembly of type IV pilus, additional pilus genes (*cupE1*), and type IV pili proteins (*pilA, pilX,* and *fimU*) were likewise upregulated. Genes associated with flagellar machinery (*flgD, flgC, flgE, flgF)* were downregulated, suggesting cells treated with R-6 have enhanced biofilm formation and adhesion, but lack motility. Additional efflux related genes such as *mexC, armR,* and *nalC* were also downregulated. Nearly the same set of anaerobic metabolism genes and regulation trends as seen in R-2E compared to DMSO were observed (*nosR, norC, norB, nosZ, coxB*). The novel addition of *coxA,* a cytochrome c oxidase subunit, was also upregulated. Meanwhile, *narG,* and *narK1* were downregulated alongside *narK2,* another nitrite transport protein. Of particular interest, a series of virulence-associated genes were repressed: *lasA* (encoding the protease LasA), *lasI* (quorum sensing circuitry), *pvdG, pvdS* (the toxin pyoverdine) and *phzF2* (the toxin phenazine).(23, 24) Finally, *phoU, phoA, pstABC, pstS,* each related to phosphate starvation, were downregulated compared to DMSO.

Interestingly, genes involved in the denitrification pathway (*narG*, *nosZ, norBCD, narK2* and the regulatory gene, *nirQ*) were downregulated at R-L6 (**S1 Table**). In alignment with this, the entire *nos* operon (*nosR, nosZ, nosD, nosF, nosY, nosL*) and most of the *nir* operon (*nirE, nirJ, nirH, nirG, nirL, PA14_06710, nirF, nirC, nirM*) were also downregulated. Like the R-6E conditions, *lasA,* along with phenazine related genes *phzA1* and *phzF1* and the pyoverdine genes, *pvdQ, pvdO,* and *pvdN,* were all repressed compared to the DMSO control. Various chemotaxis and motility related genes (*pilA, fliC, cheA, cheB, cheW, PA14_02250*) were exclusively repressed in R-6L. Lastly, pyochelin related genes (*pchABC, pchEFG, fptAB)*, MexAB-OprM related genes (*armR, nalC, mexAB*), and pel polysaccharide genes (*pelAB, pelDE,* and *pelG*) were upregulated by R-6 (**S4 Fig**). Taken together, these data show that while R-2 and R-6 are both putative HK inhibitors and are structurally similar, they cause distinct alterations in gene expression. These differences could stem from distinct HK targets leading to downstream fluctuations such as anaerobic metabolism for R-2 treatment, and siderophore and toxin genes for R-6 treatment.

### Mex Family Efflux Pump Expression

One of the most highly upregulated genes found in the R-6E and R-6L datasets was *armR,* encoding ArmR, an anti-repressor of MexR, which represses the *mexAB-oprM* operon (**Fig 2A**). Thus, upregulation of *armR* should be accompanied by upregulation of *mexAB-oprM*. We found *mexA* and *mexB* were also upregulated by R-6 (fold changes of 2.30 and 2.22, respectively, **Fig 2B, S1 Table**), as expected. Curiously, in R-2 treatments, *armR* expression was not significantly different but expression of *mexR,* and *mexAB-oprM* was repressed in R-2L (**Fig 2B, S1 Table**). Noting this contrasting effect on expression, we looked for expression changes in the additional *mex* genes. R-2 treatment resulted in increased expression of *mexEF*-*oprN* in R-2L while *mexC* was significantly downregulated in R-6E (**Fig 2B**). Given the structural similarity between R-2 and R-6, we did not expect drastic alterations in expression of different efflux pumps for each compound. Additionally, as accumulation of small molecule inhibitors is already difficult in PA14, we were interested to understand how R-2 and R-6 changed expression of efflux pumps.

To evaluate this expression change, we cloned a *lacZ* reporter downstream of the *armR* and *mexAB-oprM* promoters and performed β-galactosidase assays to report changes in expression (plasmids and primers used in these studies in **Tables S2** and **S3**). When treated with R-2 and R-6, both PA14 P*_armR_*-*lacZ* (ECA215) and PA14 P*_mexAB-oprM_-lacZ* (ECA347) revealed elevated expression as determined by β-galactosidase assays that corroborated our RNA-sequencing results (**Fig 2C**). While the response to R-6 was dose responsive, this was not the case for treatment with R-2 (**Fig 2D**) implying that R-2 may directly impact *mexEF-oprN* expression.

### Investigating Chemical Structure and Efflux Activation Relationship

We expanded our studies to a library of additional molecules to explore the relationship between chemical structure and *armR* expression. Prior work gave us an extensive library of HK inhibitors sharing the 2-aminobenzothiazole core scaffold.(24) This library, termed the B-series, is composed of 41 derivatives of the 2-aminobenzothiazole scaffold (**Fig 1A**) with modifications exclusively at the 6-position of the ring with various linkers (e.g., ether, sulfone, and sulfoxide moieties) and appended groups (e.g., pyrazoles, pyrimidines, oxazoles). PA14 P*_armR_*-lacZ was treated with all B-series inhibitors and their β-galactosidase activity measured and categorized into “high” or “low” activators of *armR* expression (**Fig 3A**). Based on our results, high and low activation was determined to be any, *β*-galactoside unit value greater than or less than 20 β-galactosidase units, respectively, with a *p*-value <0.0001 above the DMSO control (42.31 ± 4.00 β-galactoside units). These metrics were selected to narrow down structures to focus on and ensure statistically different values compared to DMSO. R-6 is a high activator (84.45 ± 0.06 units) while R-2, with a value nearly identical to DMSO (45.78 ± 0.47 units) is deemed as a “non-activator”.

**Fig 3:**
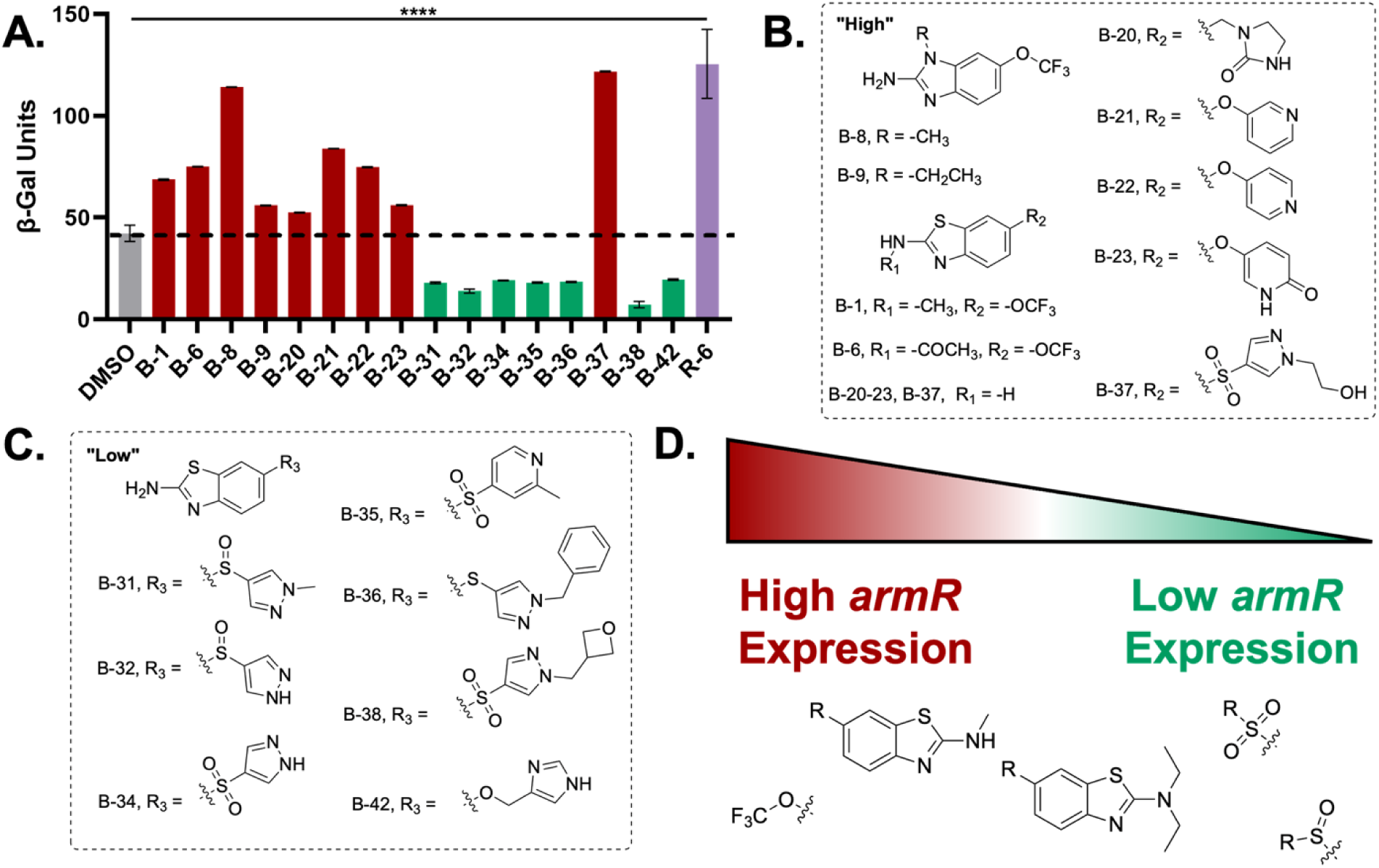
Chemical Motifs Related to *armR* Expression. A. B-series inhibitor treatments (200 µM) with PA14::P*_armR_*-lacZ strain. All, *β*-galactosidase results were compared to DMSO control, with “high” and “low” activators greater or less than two standard deviations above DMSO’s, *β*-galactoside units, respectively. Statistical significance determined by one-way ANOVA Brown-Forsythe and Welch. *p*-value ****<0.0001. All inhibitors shown have *p*-values <0.0001 compared to the DMSO control. Structures of “high” and “low” accumulating B-series inhibitors are shown in panels B and C, respectively. D. Chemical motifs that scale from high to low activation of *armR*.

Evaluation of structures within and across high, low, and non-activator categories resulted in a few trends related to specific chemical motifs. For high activation, every structure contained a -CF_3_ group, most often as -OCF_3_ (**Fig 3B**). Similarly, many structures yielding high *armR* activation included those with a small alkyl group off of the exocyclic amine in the 2-aminobenzothiazole core. Interestingly, addition of a bulky substituent to the same position seemed to lower activation. Finally, either a sulfonyl or sulfoxide linker between the 2-aminobenzothiazole core and the remainder of the inhibitor was consistently found in low *armR* activation inhibitor structures (**Fig 3C**). Combining a high and low chemical motif, such as inhibitor B-5, containing a -OCF_3_ group and significant bulk (i.e., ethyl groups) off the exocyclic amine, often resulted in the inhibitor being classified as a non-activator (**S5A Fig**). Within non-activator structures, few motifs seemed to be conserved across the group, with a notable exception of every ether linker that connected the 2-aminobenzothiazole core to any group that was not -CF_3_ was a non-activator (**S5B Fig**). When all trends are evaluated, we can start to decipher motifs that activate or do not activate efflux (**Fig 3D**).

### Synergy of Efflux Pump Deficient Strains and Virulence Inhibitors

Mex efflux pumps are best known for their role in ABR, and therefore the antibiotic substrate scope for each efflux pump is well studied. For example, imipenem is exclusively excreted from the cell by MexEF-OprN. We postulated that a synergistic effect may occur with cotreatment of R-6 and imipenem, as R-6 treatment should lower the expression level of *mexEF-oprN* and ultimately lead to decreased imipenem efflux by MexEF-OprN.(3, 31, 32) To evaluate this, we performed a checkboard assay to determine synergy between compounds. A classic checkerboard assay uses a microbroth dilution for both antibiotic and the compound tested across a multi-well plate. If growth inhibition occurs at lower concentrations than the minimum inhibitory concentration (MIC) for a compound of interest, the fractional inhibitory concentration (FIC) can be calculated. FICs with any value <0.5 represent synergy, values ranging from 0.5-4 represent no effect and values >4 demonstrate antagonism.(33) The MICs for R-6 and imipenem were 3200 µM and 4 µg/mL, respectively (**S4 Table**). As predicted, the checkerboard assay revealed an FIC of 0.20 between R-6 and imipenem, indicating a synergistic relationship (**Table 1**). Next, we sought to determine if repression of *mexAB-oprM* also increased sensitivity to antibiotics. However, no antibiotic is exclusively extruded by MexAB-OprM. (3, 31, 32) Nevertheless, we hypothesized a synergistic effect may still be seen with the known substrate, ciprofloxacin. Contrary to our expectation, the resulting FIC of 0.54 indicates there is not a synergistic relationship between R-2 and ciprofloxacin (**Table 1**). However, given that this is a borderline value, we postulated that if R-2 was able to accumulate to a higher extent within PA14, the FIC value may drop below the synergy threshold. To test this, we created in-frame deletions of each of the four main Mex efflux pumps, Δ*mexAB-oprM* (ΔAB, ECA231), Δ*mexCD-oprJ* (ΔCD, ECA343), Δ*mexEF-oprN* (ΔEF, ECA244), and Δ*mexXY* (ΔXY, ECA337) in the PA14 WT background (**Table 2**) to investigate synergy within various deletion strains. Since R-2 increases the transcripts of *mexEF-oprN*, ΔEF PA14 was selected to re-evaluate synergy between R-2 and ciprofloxacin. The FIC dropped to 0.40 with PA14 ΔEF (**Table 1**), though removal of other efflux pumps was not sufficient to cause this shift to synergy. For example, use of PA14 ΔXY, resulted in an FIC increase to 0.74 (**Table 1**). This also may suggest that R-2 is extruded through MexEF-OprN with a higher affinity than MexXY-OprM, consistent with the transcriptional profile in response to R-2L treatment.

**Table 1:** FIC Values of HK Inhibitors and Antibiotics.

| Pairing | Strain | FIC |
| --- | --- | --- |
| R-6 & Imipenem | WT | 0.20 |
| R-2 & Ciprofloxacin | WT | 0.54 |
| R-2 & Ciprofloxacin | ΔEF | 0.40 |
| R-2 & Ciprofloxacin | ΔXY | 0.75 |

**Table 2:** Strains Utilized in This Study.

| Bacterial Strain | Relevant genotype or characteristic(s) | Source |
| --- | --- | --- |

| *P. aeruginosa* strains |  |  |
| --- | --- | --- |
| ECA237 | Wild type; UCBPP-PA14 | (34) |
| ECA215 | PA14 $P_{armR}$-*lacZ* PA14 | This study |
| ECA347 | PA14 $P_{mexAB-oprM}$-*lacZ* | This study |
| ECA231 | $\Delta AB$; $\Delta mexAB-oprM$ PA14 | This study |
| ECA343 | $\Delta CD$; $\Delta mexCD-oprJ$ PA14 | This study |
| ECA244 | $\Delta EF$; $\Delta mexEF-oprN$ PA14 | This study |
| ECA337 | $\Delta XY$; $\Delta mexXY$ PA14 | This study |
| ECA307 | $\Delta^2$; $\Delta AB\Delta EF$ PA14 | This study |
| ECA338 | $\Delta^3$; $\Delta AB\Delta EF\Delta CD$ PA14 | This study |
| ECA344 | $\Delta^4$; $\Delta AB\Delta EF\Delta CD\Delta XY$ PA14 | This study |
| *E. coli* strains |  |  |
| ECA? | DH5α; High plasmid copy number | Invitrogen |
| ECA134 | SM10; Conjugation donor strain for PA14 mating | (35) |

### Impact of Inhibitor Accumulation on PA14 Motility Phenotypes

Our RNA-sequencing and synergy results show that R-2 and R-6 are likely extruded by different efflux pumps and to different extents. To evaluate contributions of the different efflux pumps, we sequentially deleted the Mex efflux pumps yielding three strains: Δ^2^ (ΔABΔEF, ‘ECA307’), Δ^3^ (ΔABΔEFΔCD, ‘ECA338’), and Δ^4^ (ΔABΔEFΔCDΔXY, ‘ECA344’) (**Table 2**). Given that transcripts relevant for both swimming and twitching were repressed by R-2L and R-6L treatment (**S1 Table**), we then evaluated whether deletion of the Mex efflux pumps would further decrease PA14 motility upon treatment with inhibitor. Interestingly, R-6 inhibited swim area to a similar extent regardless of the efflux pump deleted (**Fig 4A, S6A Fig**). Furthermore, as there was not a significant difference seen in swim area between WT or Δ^4^ PA14 upon treatment with R-6, efflux is likely not why R-6 is failing to cause potent swimming inhibition. R-2 swim inhibition results are in stark contrast to R-6, where each additional efflux pump deletion leads to more swim inhibition, ultimately abolishing swim activity in Δ^4^ PA14 (**Fig 4A**). Interestingly, removing a single efflux pump and treating with R-2 never resulted in a significant difference in swim inhibition (**S7A Fig**). At 48 hr, these trends held for R-2, but R-6 did not appear to have a reduction in swim radius in Δ^2^ and Δ^3^ PA14. These data suggest that at longer timepoints, other efflux pumps may play a role in R-6 mediated swim inhibition, or that recognition of this compound for extrusion may be slower or less efficient (**S6B Fig, S7B Fig**).

**Fig 4:**
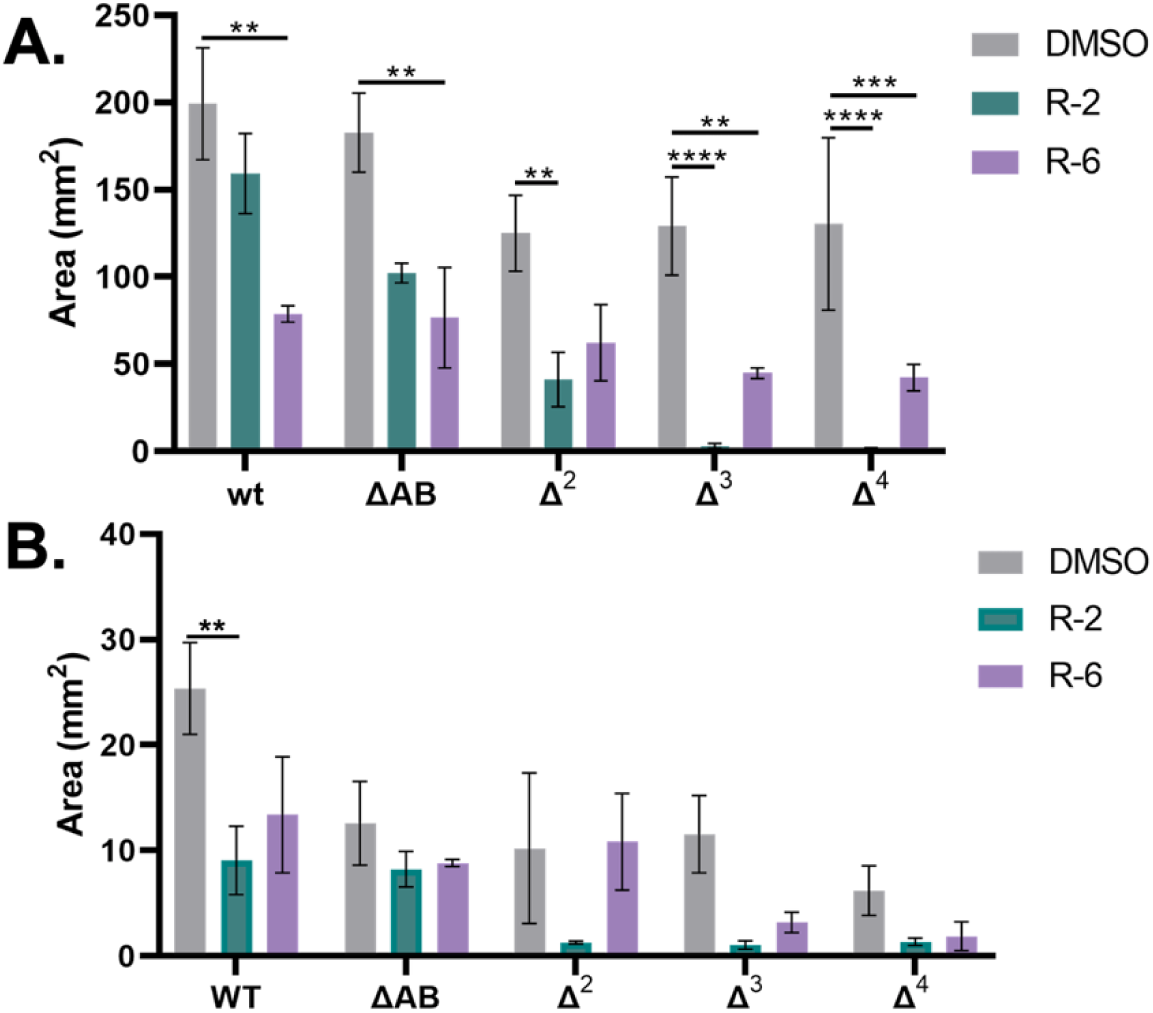
Swim and Twitch Inhibition with Efflux Pump Deletion Strains and HK Inhibitor Treatment. A. Quantified swim area for DMSO, R-2 or R-6 (200 µM) treatment at 24 hr (n=3) for WT, ΔAB, Δ^2,^ Δ^3^, and Δ^4^ PA14. B. Quantified twitch area for DMSO, R-2 or R-6 (200 µM) treatment at 24 hr (n=3) for WT, ΔAB, Δ^2,^ Δ^3^, and Δ^4^ PA14. Statistical significance determined by two-way ANOVA. *p*-value *<0.05, ** <0.005, ***<0.0005, ****<0.0001.

Twitching motility was similarly tested at 24 and 48 hr for both inhibitor treatments with each efflux pump deficient strain of PA14. Not surprisingly, observed trends were similar between the twitching data and swimming datasets. R-2 twitch inhibition increased when subsequently removing efflux pumps (**Fig 4B**). Intriguingly, for twitching there was little difference in twitch area from Δ^2^ to Δ^3^ PA14. Twitching area is small in comparison to swim area, and potent inhibition was reached with R-2 treatment in the Δ^2^ strain. Therefore, we suggest there was not sufficient area to continue to see differences in the Δ^3^ strain, rather than a lack of recognition by the MexCD-OprJ efflux pump. Indeed, at 24 hr, PA14 ΔCD exhibited twitch inhibition compared to DMSO when treated with R-2 (**S8A Fig, S9A Fig**). Interestingly, there was a trend towards higher accumulation of R-6 in Δ^3^ and Δ^4^ PA14 leading to stronger twitching inhibition. These trends held through the 48 hr timepoint (**S8B Fig, S9B Fig**). This could also be due to stronger target engagement and therefore a consequent stronger phenotype compared to the swim data. For example, *pilA*, the major subunit of the type IV pilus, was downregulated the most after treatment with R-6 during log phase (R-6L) (**S1 Table**). Meanwhile, *flrA, flrC* and *fliA* are major regulators of the flagellar assembly required for swimming motility. In R-2L, each of these were significantly downregulated, but there was no significant change in these transcripts in R-6L, possibly explaining why more potent inhibition was observed for swimming motility under R-2 treatment.

### TCSs Connection to Transcriptomic Alterations and Inhibition of Virulence Phenotypes

Within the R-2 datasets, *mexEF-oprN* genes and those within its regulon were highly expressed. The regulation of MexEF-OprN hinges on *mexS,* which is sensitive to the oxidative changes caused by quinolone compounds and then activates *mexT* increasing *mexEF-oprN* transcription.(26, 27) The R-2E dataset was dominated by redox sensitive enzymes, possibly tied to the oxidative switch that *mexS* is known to respond to. Interestingly, the TCS CzcSR, linked to Zn^2+^ homeostasis and carbapenem resistance, was also significantly upregulated.(36) Deletion of the response regulator CzcR caused complete loss of swimming in *P. aeruginosa* despite activation of CzcS. (37) If CzcS is active, but R-2 inhibits it, we would expect our results to phenocopy the CzcR deletion in the swimming assay, and thus show complete swimming inhibition. This matches our R-2L dataset, especially under high accumulating conditions, such as in treatment of the Δ^4^ strain (**Fig 4A**). Furthermore, CzcR regulates expression of the global zinc regular *zur* (formally *np20*) as well as the *znuB* and *znuC* genes, which are decreased more than two-fold further suggesting repression of this TCS in R-2 treatments. Unique to R-6 treatments (**S10 Fig**), but minimal perturbations seen for R-2 (**S11 Fig**), were phenazine related genes, explaining our prior observed phenotypic response of more potent pyocyanin reduction by R-6 treatment.(24)

Martínez and coworkers previously found that hyperactivity of Mex efflux pumps caused a drop in the intracellular pH as efflux pumps are proton antiporters. Acidification of the cytosol led *P. aeruginosa* to further shift its metabolism to anaerobic pathways relieving the stress of low pH in the cell. (38, 39) Our data show that both treatments yielded increased expression of both efflux pumps and the denitrification pathway, suggesting an overall shift in anaerobic metabolism (**S12-S15 Figs**). We posit that a similar phenotype occurs for R-6, as the highest increase in efflux pump expression was observed under this condition. However, it is worth noting that the *narG* and *narK2* operon (*narK1K2GHJI*) shares a promoter region with genes encoding the TCS NarXL, responsible for nitrate sensing. Furthermore, NarX is known to regulate the *nar* and *nir* operon, as well as genes encoding other major nitrate respiration regulators, NirQ and Dnr.(40) We observed repression in the *nar* and *nir* operons in all timepoints, and *nirQ* in R-6L implicating repression of NarXL. R-6 treatment *of P. aeruginosa* resulted in high levels of siderophore gene expression and repression of phosphate-related genes (**Fig 5B**), possible downstream responses linked to the shifting nitrate metabolism. In R-2, many other nitrosative related genes were perturbed and we hypothesize this could cause increased *nirS* and *norBC* expression with a decrease in *nosZ* and *narGHI*. So, while both inhibitors cause perturbed denitrification, we it is likely they act through distinct mechanisms. Finally, across all treatments, various flagellar assembly related genes were repressed (**S16-S19 Figs**). This aligns well with our swimming phenotypic results (**Fig 4A)** and prior swarming repression.(23, 24)

**Fig 5:**
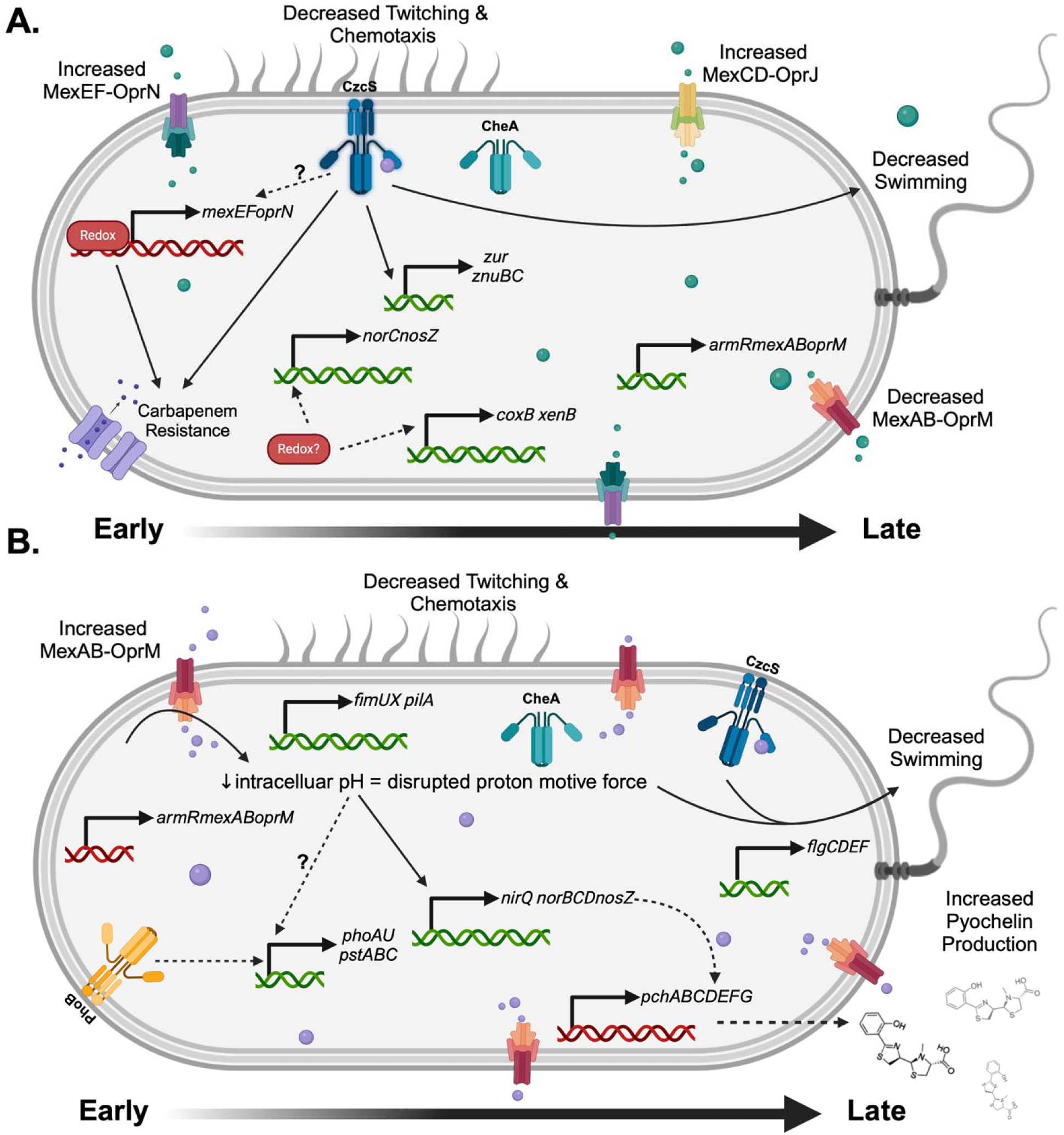
Summary of Transcriptomic and Phenotypic Response to R-2 and R-6 Treatment. Overview of R-2 (A) and R-6 (B) treatment from early to late timepoints in RNA-seq dataset. Green genes indicate downregulation whereas red indicate upregulation. Black arrows indicate known relationships between genes, proteins and/or phenotypes. Dashed arrows indicate postulated relationships between genes, proteins and/or phenotypes.

## Conclusion

Overall, this investigation into the transcriptomic response of *P. aeruginosa* enabled us to discover the expression differences likely accounting for our contrasting phenotypic responses of R-2 and R-6. Furthermore, it clearly showcased the drastic differences that similar chemical structures can cause in a cell. While trends from our small 2-aminobenzothiazole library were drawn for efflux activation, the efflux pump-deficient strains generated through this work will enable us to continue defining molecular motifs that activate different Mex efflux pumps. Additionally, these efflux-pump deficient strains can be leveraged to reach sufficient intracellular concentrations of R-2 and R-6 that could enable discovery of novel cellular targets. Our transcriptomic results indicate entire TCSs are perturbed by treatment with R-2 or R-6 (**S20 and S21 Figs**), and thus should complement cellular targeting studies.

## Materials and Methods

### Materials

All chemicals were obtained through the following companies. LB Media (Sigma, powder, cat #L3022), Agar (Sigma, powder, cat #A1296), Plastic 17 x100 mm Culture tubes (25 pack) (VWR, cat#60818-703), Breathe-Easy sealing membrane (Sigma Aldrich, cat# Z380059) RNA Extraction Mini Kit (Qiagen, Cat # 74104) RNAprotect Reagent (Qiagen, cat# 76506), RNase Spray (ThermoFisher, cat# 21-236-21), Invitrogen TE Buffer RNase free (ThermoFisher, Cat #AM9849), Invitrogen TURBO DNase (2U/uL) (ThermoFisher, cat #AM2238), Gel loading dye, purple (New England Biolabs, cat# B7024A), TriDye 1kb DNA Ladder (New England Biolabs, cat# N3272S), GelRed (Thermofisher, SCT123), 96-Well PCR Plates, high profile, semi-skirted (BioRad, cat#2239441), Microseal ‘B’ PCR Plate Sealing Film, adhesive, optical (BioRad, cat#MSB1001), *E. coli* DH5⍰ (Invitrogen, cat # 12297-016).

### Growth Conditions

The WT PA14, PA14 efflux pump deletions, and *lacZ* fusions were streaked onto Lennox Broth (LB) agar plates from a frozen glycerol stock and grown overnight at 37 °C. From these plates, a single colony was inoculated into fresh LB media (1 mL) and grown overnight at 37 °C, shaking at 220 RPM. A 1:100 dilution of the overnight culture was performed in fresh LB media and grown until desired OD_600_ (Genesys 30 Visible Spectrophotometer). Cells were then treated in accordance with each assay.

### RNA Extraction

PA14 diluted from overnight cultures were diluted into fresh LB media containing DMSO, R-2 or R-6 (200 µM) and grown to OD_600_ ∼0.9 or ∼1.2. Four biological replicates were grown for every treatment condition. For each sample, 2X volume of RNA protect reagent (Qiagen) was added, cells vortexed and incubated at room temperature for 10 min. Following incubation, 1 mL aliquots of treated cells were spun down (10 min, 5000g), supernatant decanted and cell pellet snap frozen at −80 °C. RNeasy Mini Kit (Qiagen) was utilized to extract RNA according to the manufacturer’s protocol combined with an additional off-column DNase treatment (Turbo DNase, Invitrogen). RNA integrity and contamination was evaluated by agarose gels after DNase treatment. RNA concentration was measured via a Tecan Spark Nanoquant plate.

### RNA-Sequencing

RNA integrity was determined by the University of Minnesota Genomics Center and samples with RNA-Integrity Numbers > 8.0 were used for sequencing. Sequencing was completed with the Illumina NovaSeq platform SP 50PE with ∼15.6M reads/sample. Quality control, data alignment and gene quantification was analyzed using the CHURP pipeline(41) at the University of Minnesota Supercomputing Institute (MSI). 2 x 50bp FASTQ paired-end reads for 24 samples (13.4 million reads average per sample) were trimmed using Trimmomatic (v0.33) enabled with the optional “-q” option; 3bp sliding-window trimming from 3’ end requiring minimum Q30. Quality control on raw sequence data for each sample was performed with FastQC. Read mapping was performed via HISAT2(v2.1.0) using the *Pseudomonas aeruginosa* genome (***ASM1462v1***) as a reference. Gene quantification was performed via Feature Counts for raw read counts. Using raw read counts, differentially expressed genes were identified using the edgeR (negative binomial, R programming) feature in CLCGWB (Qiagen, Redwood City, CA). We filtered the generated list based on a minimum 2X absolute fold change, and FDR corrected p < 0.05.

### Synergy Assay

Antibiotic stocks were prepared in sterile MilliQ and stored at −20 □ except for imipenem which was made fresh the day of each experiment. R-2 and R-6 stocks were stored in DMSO at −20 □. For each antibiotic a 4X stock was generated in LB media from the desired final concentration in column one of the 96-well plate. In column one 100 µL of the 4X antibiotic stock was dispensed. In all other columns 50 µL of LB media was dispensed. Antibiotic from column one was serially diluted two-fold across columns two-eleven. Similarly, a 4X stock of R-2 or R-6 was generated in LB media (final DMSO < 2% v/v) from the desired final concentration in column A of the 96-well plate. This was serially diluted two-fold in sterile microcentrifuge tubes. Once complete, 50 µL of the 4X stock was placed in row A, followed by the next dilution in row B, and so on until row G. LB media (50 µL) was added to row H and column 12 to serve as internal MIC controls. PA14 strains of interest were diluted from overnight cultures 1:100 into fresh LB. Diluted cells (100 µL) were added to every well. Plates were sealed with Breathe-Easy (Sigma Aldrich) plate films and incubated at 37 □ for ∼16hr. OD_600_ measurements were taken on a Tecan Spark. FIC values were calculated with the following equation where MIC_A_ is the MIC of the antibiotic, MIC_B_ is the MIC of either R-2 or R-6 (MIC of R-2 was set at 2000 µM as this was the solubility limit in LB media) and MIC_C_ is the combined MIC of antibiotic and inhibitor.

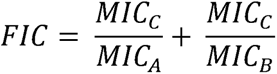

### Swimming Assay

Procedure from Ha et al.(42) Briefly, 0.3% agar plates containing M8 media (M8 salts, 0.2% w/v glucose, 0.5% w/v casamino acids, 1 mM MgSO_4_) containing DMSO, R-2 or R-6 (200 µM) were poured (∼25-30 mL) into a sterile petri dish (Corning, 25 x100 mm) the day of an experiment and allowed to cool to room temperature. Overnight cultures of PA14 were transferred to sterile microcentrifuge tubes. Sterile toothpicks were dipped into overnight cultures, ensuring no excess culture on the toothpick, and rapidly stabbed directly into the middle of the agar plate. Plates were incubated for up to 48 hr at 37 □ upright.

### Twitching Assay

Procedure from Turnbull and Whitchurch.(42) Briefly, 1.5% LB agar plates containing DMSO, R-2 or R-6 (200 µM) were poured (∼10 mL) and swirled across the bottom of a sterile petri dish (Corning, 25 x 100 mm) the day of an experiment and allowed to cool to room temperature. Overnight cultures of PA14 were transferred to sterile microcentrifuge tubes. Sterile toothpicks were dipped into overnight cultures, ensuring no excess culture on the toothpick, and rapidly stabbed directly into the bottom of the agar plate between the plastic and agar. Plates were incubated for up to 48 hr at 37 □ upright.

## Supporting information

Lembke Supporting Information

## Supplemental Information

Strain generation, β-galactosidase assays, swimming and twitching quantification, additional experimental details, materials, and methods can be found in the supplemental information.

## Acknowledgements

The authors acknowledge the Minnesota Supercomputing Institute (MSI) at the University of Minnesota for providing resources that contributed to the results reported within this paper. Additionally, we thank the University of Minnesota Genomics Center (RRID: SCR_012413) for completing the QC, library generation and sequencing for our RNA-Sequencing experiment, as well as their technical support. We would like to thank Dr. Juan E. Abrahante Lloréns for his expertise in bioinformatics and processing our RNA-Sequencing data. Whole genome sequencing for each deletion strain was completed by Seqcenter (Philadelphia, PA). Lastly, we acknowledge BioRender for help in our figure generation.

## Authors Contribution Statement

Hannah Lembke: Ran all experiments for manuscript. Generated some deletion strains (CD, XY, triple and quadruple deletions). Wrote main manuscript and SI, figure generation for main manuscript and SI.

Kelsie Nauta: Contributed to project development, generated some of the knockout strains and *lacZ* reporters as well as bacterial genetics methods.

Ryan Hunter: Project oversight, conception and design of experiments, RNAseq analysis, manuscript editing, funding acquisition.

Erin Carlson: Project oversight, conception and design of experiments, manuscript editing, funding acquisition.

---

**Suggested TOC:**
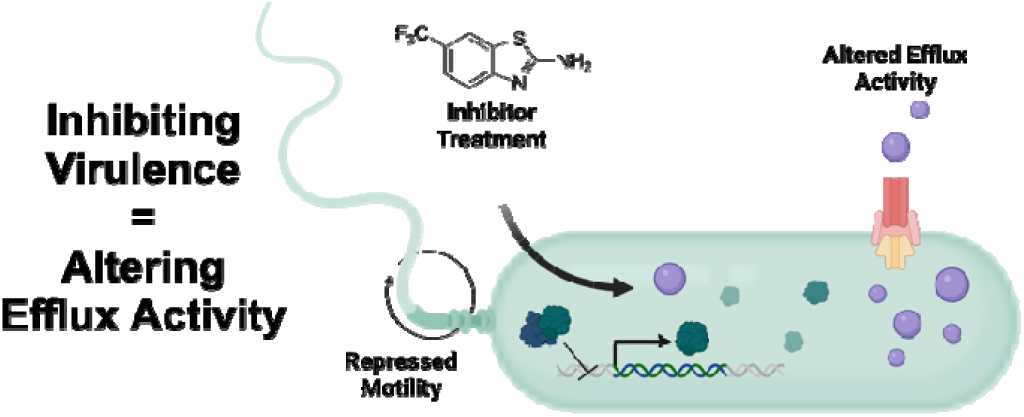

## References

1. Rasko DA, Sperandio V. Anti-virulence strategies to combat bacteria-mediated disease. Nature Reviews Drug Discovery. 2010;9(2):117–28.

2. Vasil ML. Pseudomonas aeruginosa. Biology, mechanisms of virulence, epidemiology. The Journal of Pediatrics. 1986;108:800–5.

3. Lister PD, Wolter DJ, Hanson ND. Antibacterial-resistant Pseudomonas aeruginosa: clinical impact and complex regulation of chromosomally encoded resistance mechanisms. Clin Microbiol Rev. 2009;22(4):582–610.

4. Chatterjee M, Anju CP, Biswas L, Anil Kumar V, Gopi Mohan C, Biswas R. Antibiotic resistance in Pseudomonas aeruginosa and alternative therapeutic options. Int J Med Microbiol. 2016;306(1):48–58.

5. Pang Z, Raudonis R, Glick BR, Lin TJ, Cheng Z. Antibiotic resistance in Pseudomonas aeruginosa: mechanisms and alternative therapeutic strategies. Biotechnol Adv. 2019;37(1):177–92.

6. Silhavy TJ, Kahne D, Walker S. The bacterial cell envelope. Cold Spring Harb Perspect Biol. 2010;2(5):a000414–a.

7. Ude J, Tripathi V, Buyck JM, Soderholm S, Cunrath O, Fanous J, et al. Outer membrane permeability: Antimicrobials and diverse nutrients bypass porins in Pseudomonas aeruginosa. Proc Natl Acad Sci U S A. 2021;118(31).

8. Geddes EJ, Gugger MK, Garcia A, Chavez MG, Lee MR, Perlmutter SJ, et al. Porin-independent accumulation in Pseudomonas enables antibiotic discovery. Nature. 2023;624(7990):145–53.

9. Dreier J, Ruggerone P. Interaction of antibacterial compounds with RND e ffl ux pumps in Pseudomonas aeruginosa. Front Microbiol. 2015;6:660.

10. Masuda N, Sakagawa E, Tsujimoto H, Ohya S, Gotoh N, Nishino T. Substrate Specificities of MexAB-OprM, MexCD-OprJ, and MexXY-OprM Efflux Pumps in Pseudomonas aeruginosa. Antimicrob Agents Chemother. 2000;44:3322–7.

11. Gervasoni S, Malloci G, Bosin A, Vargiu AV, Zgurskaya HI, Ruggerone P. Recognition of quinolone antibiotics by the multidrug efflux transporter MexB of Pseudomonas aeruginosa. Phys Chem Chem Phys. 2022;24(27):16566–75.

12. Lopez CA, Travers T, Pos KM, Zgurskaya HI, Gnanakaran S. Dynamics of Intact MexAB-OprM Efflux Pump: Focusing on the MexA-OprM Interface. Sci Rep. 2017;7(1):16521.

13. Silvia Gervasoni, Jitender Mehla, Olga Lomovskaya, Charles R. Bergen, Inga V. Leus, Enrico Margiotta, et al. Molecular determinants of avoidance and inhibition of Pseudomonas aeruginosa MexB efflux pump. mBio. 2023.

14. Lau GW, Hassett DJ, Ran H, Kong F. The role of pyocyanin in Pseudomonas aeruginosa infection. Trends Mol Med. 2004;10(12):599–606.

15. Van Alst NE, Picardo KF, Iglewski BH, Haidaris CG. Nitrate sensing and metabolism modulate motility, biofilm formation, and virulence in Pseudomonas aeruginosa. Infect Immun. 2007;75(8):3780–90.

16. Yashkin A, Rayo J, Grimm L, Welch M, Meijler MM. Short-chain reactive probes as tools to unravel the Pseudomonas aeruginosa quorum sensing regulon. Chem Sci. 2021;12(12):4570–81.

17. Qin S, Xiao W, Zhou C, Pu Q, Deng X, Lan L, et al. Pseudomonas aeruginosa: pathogenesis, virulence factors, antibiotic resistance, interaction with host, technology advances and emerging therapeutics. Signal Transduct Target Ther. 2022;7(1):199.

18. Tiwari S, Jamal SB, Hassan SS, Carvalho PVSD, Almeida S, Barh D, et al. Two-Component Signal Transduction Systems of Pathogenic Bacteria As Targets for Antimicrobial Therapy: An Overview. Front Microbiol. 2017;8:1878-.

19. Sultan M, Arya R, Kim KK. Roles of Two-Component Systems in Pseudomonas aeruginosa Virulence. Int J Mol Sci. 2021;22(22).

20. Francis VI, Stevenson EC, Porter SL. Two-component systems required for virulence in Pseudomonas aeruginosa. FEMS Microbiol Lett. 2017;364(11).

21. Stock AM, Robinson VL, Goudreau PL. Two Component Signal Transduction. Annu Rev Biochem 2000;69:183–215.

22. Wilke KE, Francis S, Carlson EE. Inactivation of multiple bacterial histidine kinases by targeting the ATP-binding domain. ACS Chem Biol. 2015;10(1):328–35.

23. Goswami M, Espinasse A, Carlson EE. Disarming the virulence arsenal of Pseudomonas aeruginosa by blocking two-component system signaling. Chemical Science. 2018;9(37):7332–7.

24. Fihn CA, Lembke HK, Gaulin J, Bouchard P, Villarreal AR, Penningroth MR, et al. Evaluation of expanded 2-aminobenzothiazole library as inhibitors of a model histidine kinase and virulence suppressors in Pseudomonas aeruginosa. Bioorganic Chemistry. 2024;153.

25. Winsor GL, Griffiths EJ, Lo R, Dhillon BK, Shay JA, Brinkman FS. Enhanced annotations and features for comparing thousands of Pseudomonas genomes in the Pseudomonas genome database. Nucleic Acids Res. 2016;44(D1):D646–53.

26. Fargier E, Mac Aogain M, Mooij MJ, Woods DF, Morrissey JP, Dobson AD, et al. MexT functions as a redox-responsive regulator modulating disulfide stress resistance in Pseudomonas aeruginosa. Journal of bacteriology. 2012;194(13):3502–11.

27. Fetar H, Gilmour C, Klinoski R, Daigle DM, Dean CR, Poole K. mexEF-oprN multidrug efflux operon of Pseudomonas aeruginosa: regulation by the MexT activator in response to nitrosative stress and chloramphenicol. Antimicrob Agents Chemother. 2011;55(2):508–14.

28. Vaillancourt M, Limsuwannarot SP, Bresee C, Poopalarajah R, Jorth P. Pseudomonas aeruginosa mexR and mexEF Antibiotic Efflux Pump Variants Exhibit Increased Virulence. Antibiotics (Basel). 2021;10(10).

29. Mima T, Sekiya H, Mizushima T, Kuroda T, Tsuchiya T. Gene cloning and properties of the RND-type multidrug efflux pumps MexPQ-OpmE and MexMN-OprM from Pseudomonas aeruginosa. Microbiol Immunol. 2005;49(11):999–1002.

30. Laverty G, Gorman SP, Gilmore BF. Biomolecular Mechanisms of Pseudomonas aeruginosa and Escherichia coli Biofilm Formation. Pathogens. 2014;3(3):596–632.

31. Piddock LJ. Clinically relevant chromosomally encoded multidrug resistance efflux pumps in bacteria. Clin Microbiol Rev. 2006;19(2):382–402.

32. Zwama M, Nishino K. Ever-Adapting RND Efflux Pumps in Gram-Negative Multidrug-Resistant Pathogens: A Race against Time. Antibiotics (Basel). 2021;10(7).

33. Hsieh MH, Yu CM, Yu VL, Chow JW. Synergy Assessed by Checkerboard A Critical Analysis. Diagn Microbiol Infect Dis. 1993;16:343–9.

34. Rahme LG, Stevens EJ, Wolfort SF, Shao J, Tompkins RG, Ausubel FM. Common Virulence Factors for Bacterial Pathogenicity in Plants and Animals. Science. 1995;268(5219):1899–902.

35. Simon R, Priefer U, Pühler A. A Broad Host Range Mobilization System for In Vivo Genetic Engineering: Transposon Mutagenesis in Gram Negative Bacteria. Bio/Technology. 1983;1(9):784–91.

36. Perron K, Caille O, Rossier C, van Delden C, Dumas J-L, Köhler T. CzcR-CzcS, a Two-component System Involved in Heavy Metal and Carbapenem Resistance in Pseudomonas aeruginosa. Journal of Biological Chemistry. 2004;279(10):8761–8.

37. Zhiqing L., Zirui X., Shuzhen C., Jiahui H., Ting L., Cheng D., et al. CzcR Is Essential for Swimming Motility in Pseudomonas aeruginosa during Zinc Stress. 2022;10(6).

38. Olivares J, Alvarez-Ortega C, Linares JF, Rojo F, Kohler T, Martinez JL. Overproduction of the multidrug efflux pump MexEF-OprN does not impair Pseudomonas aeruginosa fitness in competition tests, but produces specific changes in bacterial regulatory networks. Environ Microbiol. 2012;14(8):1968–81.

39. Olivares J, Alvarez-Ortega C, Martinez JL. Metabolic compensation of fitness costs associated with overexpression of the multidrug efflux pump MexEF-OprN in Pseudomonas aeruginosa. Antimicrob Agents Chemother. 2014;58(7):3904–13.

40. Cheung J, Hendrickson WA. Structural analysis of ligand stimulation of the histidine kinase NarX. Structure. 2009;17(2):190–201.

41. Baller J, Kono T, Herman A, Zhang Y. Churp. Proceedings of the Practice and Experience in Advanced Research Computing on Rise of the Machines (learning)2019. p. 1–5.

42. Pseudomonas Methods and Protocols. Filloux A, Ramos J-L, editors. New York: Springer; 2014.

