## Supplementary material for "Evaluating the Link Between Efflux Pump Expression and Motility Phenotypes in *Pseudomonas aeruginosa* Treated with Virulence Inhibitors": Lembke Supporting Information

**<u>Experimental Methods</u>**

Materials and Methods

Plasmid Generation

*E. coli* Transformation

*P. aeruginosa* Conjugation

*P. aeruginosa* Counter Selection

β-Galactosidase Assay

Quantifying swimming and twitching in ImageJ

**<u>Supporting Figures, and Tables</u>**

**S1 Table:** Differentially Expressed Genes in R-2 and R-6 Treated Samples.

**S2 Table:** Plasmids used in this study

**S3 Table:** Cloning primers used in this study

**S1 Fig:** R-2E STRING Analysis

**S2 Fig:** R-2L STRING Analysis

**S3 Fig:** R-6E STRING Analysis

**S4 Fig:** R-6L STRING Analysis

**S5 Fig:** B-series Non-Hits Structures and β-Gal Values

**S4 Table:** MIC Values

**S6 Fig:** Swimming Images

**S7 Fig:** Quantified Swimming Area

**S8 Fig:** Twitching Images

**S9 Fig:** Quantified Twitching Area

**S10 Fig:** Pathview Pathway of R-6 Phenazine Biosynthesis

**S11 Fig:** Pathview Pathway of R-2 Phenazine Biosynthesis

**S12 Fig:** Pathview Pathway of R-2E Nitrate Respiration

**S13 Fig:** Pathview Pathway of R-2L Nitrate Respiration

**S14 Fig:** Pathview Pathway of R-6E Nitrate Respiration

**S15 Fig:** Pathview Pathway of R-6L Nitrate Respiration

**S16 Fig:** Pathview Pathway of R-2E Flagellar Assembly

**S17 Fig:** Pathview Pathway of R-2L Flagellar Assembly

**S18 Fig:** Pathview Pathway of R-6E Flagellar Assembly

**S19 Fig:** Pathview Pathway of R-6L Flagellar Assembly

**S20 Fig:** Pathview Pathway of R-2 Two-Component Systems

**S21 Fig:** Pathview Pathway of R-6 Two-Component Systems

**Materials & Methods**

*Materials.* All chemicals were obtained through the following companies. LB Media (Sigma, powder, cat #L3022), Agar (Sigma, powder, cat #A1296), Plastic 17 x100 mm Culture tubes (25 pack) (VWR, cat#60818-703), Breathe-Easy sealing membrane (Sigma Aldrich, # Z380059) RNA Extraction Mini Kit (Qiagen, Cat # 74104) RNAprotect Reagent (Qiagen, Cat# 76506), RNase Spray (ThermoFisher, Cat# 21-236-21), Invitrogen TE Buffer RNase free (ThermoFisher, Cat #AM9849), Invitrogen TURBO DNase (2U/uL) (ThermoFisher, Cat # AM2238), Gel loading dye, purple (New England Biolabs, cat# B7024A), TriDye 1kb DNA Ladder (New England Biolabs, Cat# N3272S), GelRed (Thermofisher, SCT123), 96-Well PCR Plates, high profile, semi-skirted (BioRad, cat#2239441), Microseal 'B' PCR Plate Sealing Film, adhesive, optical ( BioRad, cat#MSB1001), Plasmid miniprep kit (Monarch, cat # T1010S).

*In-gel fluorescence detection.* After agarose electrophoresis, gels were scanned on a GE Typhoon FLA 9500 gel scanner in the EtBr channel (λex/λem 525 nm/600 nm).

*Strain and plasmid construction.* All plasmids used in cloning are listed in Table S2, and the respective primers used can be found in Table S3. All primers were synthesized by Integrated DNA Technologies (Coralville, IA). Plasmids were constructed by isothermal assembly.(1) Regions of plasmids constructed using PCR were verified by DNA electrophoresis. All plasmids were propagated using *E. coli* DH5α as the cloning host. To construct lacZ fusions, 1 kb of DNA upstream of the gene of interest was cloned into the pMini-CTX-lacZ plasmid, digested with BamHI and EcoRI, and selected for with 100 µg/mL carbenicillin (carb), using the primers in **S3 Table**. To construct deletion mutants, we cloned 1 kb of DNA upstream and 1 kb downstream of the site of the desired deletion using primers listed in **S2 Table** into the pEXG2tc plasmid digested with EcoRI and HindIII and selected for with 10 µg/mL tetracycline (tet) and 10% sucrose within the multicloning site. Mutants were constructed by shifting temperatures as previously described.(1) Plasmids were extracted and purified with a plasmid mini prep kit (New England Biolabs), and confirmed via sequencing (ACGT).

*E. coli transformation.* Plasmids were introduced into chemically competent *E. coli* DH5α(2) post ITA or *E. coli* SM10 by heat shock. Briefly, recipient cells were thawed on ice. Plasmid DNA (5 µL) was resuspended with cells and mixed thoroughly by pipetting. Cells incubated on ice for 10 min followed by heat shock at 42 °C for 45 sec. Cells were immediately resuspended in LB media (500 µL) and incubated at 37 °C for 1 hr. Cells (100 µL) were then plated on selection agar plates, tet (10 µg/mL for pEXG2tc vectors and carb 100 µg/mL for pMiniCTX-lacZ vectors), with sterile glass beads and incubated overnight at 37 °C. Single colonies were restreaked on selection plates and plasmid DNA extracted via a mini kit (New England Biolabs), digested and confirmed via agarose electrophoresis.

*P. aeruginosa Conjugation.* Recipient *P. aeruginosa* strain and *E. coli* SM10 mating strain were grown overnight in LB media or LB media with selection respectively at 37 °C, 220RPM. The recipient and donor strains were mixed at four ratios (1:1, 1:3, 1:4, 1:5) and underwent heat shock (42 °C, 10 min), spun down (8000*g*, 2 min) resuspended in LB media (60 µL) and plated on LB agar, incubated overnight at 30 °C. Transconjugants were isolated by plating on Vogel-Boner Minimal Media (VBMM) (10X recipe: 2g $MgSO_4 \cdot 7H_2O$, 20g Citric Acid, 35g $NaNH_4HPO_4 \cdot$

$4H_2O$, 100g $K_2HPO_4$, 1L MilliQ, pH=7) plates (1x VBMM, 0.1 % Casamino Acids, 1 mM $MgSO_4$) containing tet.

*P. aeruginosa Counter Selection.* All transconjugants that grew on VBMM selection plates were restreaked onto LB no salt, 10% sucrose plates (1.0% tryptone, 0.5% yeast extract, 10.0% sucrose) and grown overnight at 30 °C. Positive deletion colonies were checked for either carb or tet sensitivity on VBMM plates and deletions confirmed through single-colony PCR. Clones were then whole genome sequenced (SeqCenter). For *lacZ* fusions all carb$^R$ clones were conjugated with the SM10 strain carrying the pFLP2 plasmid for removal of the antibiotic cassette followed by sucrose counter selection.

*β-Galactosidase assays.* Overnight cultures were subcultured 1:100 in fresh LB containing DMSO, R-2 or R-6 (200 μM) and incubated to mid-log phase (OD of 0.8 to 1.5) at 37 °C, 200 RPM. One mL of each subculture was pelleted (8,000*g*, 5 min) and resuspended in one ml of Z-buffer (0.06 M $Na_2HPO_4$, 0.04 M $NaH_2PO_4*H_2O$, 0.01 M KCl, 0.001 M $MgSO_4$). Cells were permeabilized by addition of 16 μl chloroform. Permeabilized cells (50 μl) were mixed with 100 μl of Z-buffer and 50 μl of 5 mg/ml ortho-Nitrophenyl-β-galactoside (ONPG) (50 μl). The $OD_{420}$ was measured over time using a Tecan Spark plate reader. β-Galactosidase activity units [(μmol of ONPG formed per minute) × $10^3$/ ($OD_{600}$ × mL of cell suspension)] were calculated as previously described.(3) Experiments were performed in technical and biological triplicate, with the means and standard deviations shown.

*Quantifying swimming and twitching in ImageJ.* For every image opened in ImageJ (NIH) the area of the petri dish was measured at least three times and averaged. The pixel count was then set to the appropriate scale for a 15x100 mm dish. The scale was kept globally throughout each picture measurement. Both area and radii were measured for each biological replicate of twitching or swimming plates. Radii were measured with the line tool at least eight times from the center of the twitch or swim (where the agar plate was stabbed) to the edge of the twitch or swim. Area was measured with the circular or shape tool for swims and twitches respectively

**S1 Table: Differentially Expressed Genes in R-2 and R-6 Treated Samples.**

| Gene | Gene ID | Annotated Function | Fold Change |
|---|---|---|---|
| | | **R-2E** | |
| | PA14_70910 | 16S ribosomal RNA | 10.18 |
| | PA14_47640 | major facilitator transporter | 9.64 |
| | PA14_28410 | hypothetical protein | 8.25 |
| | PA14_16660 | metal transporting P type ATPase | 6.83 |
| | PA14_39420 | PA I galactophilic lectin | 6.69 |
| *norC* | PA14_06810 | nitric oxide reductase subunit C | 6.41 |
| *norB* | PA14_06830 | nitric oxide reductase subunit B | 5.53 |
| | PA14_55631 | 23S ribosomal RNA | 5.44 |
| | PA14_06800 | hypothetical protein | 5.15 |
| | PA14_47610 | transcriptional regulator | 5.03 |
| | PA14_39060 | lipoprotein | 4.78 |
| | PA14_64530 | hypothetical protein | 4.73 |
| *pyeM* | PA14_56640 | MFS transporter | 4.29 |
| | PA14_22420 | hypothetical protein | 4.25 |
| *nirO* | PA14_06790 | cytochrome c oxidase subunit | 4.20 |
| | PA14_67540 | hypothetical protein | 4.10 |
| *nosR* | PA14_20230 | regulatory protein NosR | 3.99 |
| | PA14_16310 | MFS permease | 3.89 |
| *mexF* | PA14_32390 | RND multidrug efflux transporter MexF | 3.75 |
| | PA14_51050 | aldehyde dehydrogenase | 3.75 |
| *nirS* | PA14_06750 | nitrite reductase | 3.47 |
| *mexE* | PA14_32400 | RND multidrug efflux membrane fusion protein MexE | 3.44 |
| | PA14_06860 | hypothetical protein | 2.95 |
| *pvdS* | PA14_33260 | extracytoplasmic function sigma 70 factor | 2.79 |
| *norD* | PA14_06840 | 8enitrification protein NorD | 2.78 |
| | PA14_49050 | hypothetical protein | 2.70 |

| | | | |
|---|---|---|---|
| *nirM* | PA14_06740 | cytochrome c551 | 2.67 |
| | PA14_35160 | phenazine utilizing monooxygenase A | 2.66 |
| *nosZ* | PA14_20200 | nitrous oxide reductase | 2.51 |
| *nirC* | PA14_06730 | c type cytochrome | 2.47 |
| *czcS* | PA14_31950 | Zn responsive histidine kinase | 2.46 |
| | PA14_62060 | 23S ribosomal RNA | 2.45 |
| | PA14_29030 | FMN oxidoreductase | 2.36 |
| *xenB* | PA14_56660 | xenobiotic reductase | 2.32 |
| *ibpA* | PA14_23680 | heat shock protein IbpA | 2.31 |
| *phnD* | PA14_20320 | phosphonate ABC transporter substrate binding protein | 2.31 |
| | PA14_39750 | amino acid permease | 2.28 |
| | PA14_26810 | two component sensor | 2.26 |
| *hslV* | PA14_66770 | ATP dependent protease peptidase subunit | 2.15 |
| *czcR* | PA14_31960 | two component response regulator | 2.14 |
| | PA14_68120 | outer membrane protein | 2.14 |
| | PA14_70860 | hypothetical protein | 2.14 |
| | PA14_49690 | oxidoreductase | 2.12 |
| *phnE* | PA14_20330 | phosphonate ABC transporter permease | 2.11 |
| *coxB* | PA14_01290 | cytochrome c oxidase subunit II | 2.09 |
| | PA14_68110 | transcriptional regulator | 2.08 |
| | PA14_46900 | hypothetical protein | 2.08 |
| *ergA* | PA14_02740 | class II aldolase/adducin domain containing protein | 2.07 |
| | PA14_23530 | secretion_protein | 2.02 |
| *narK2* | PA14_13770 | nitrite extrusion protein 2 | -2.01 |
| *mdcC* | PA14_02570 | malonate decarboxylase subunit delta | -2.02 |
| *narG* | PA14_13780 | Respiratory nitrate reductase alpha subuit | -2.03 |
| | PA14_19350 | hypothetical protein | -2.03 |
| *pa1L* | PA14_31290 | PA I galactophilic lectin | -2.05 |
| | PA14_11320 | hypothetical protein | -2.06 |

| | | | |
|---|---|---|---|
| | PA14_55110 | hypothetical protein | -2.08 |
| *np20* | PA14_72560 | transcriptional regulator np20 | -2.13 |
| | PA14_41290 | hypothetical protein | -2.13 |
| *tonB2* | PA14_02490 | tonB2 hypothetical protein | -2.20 |
| *znuC* | PA14_72580 | zinc transporter | -2.20 |
| *znuB* | PA14_72590 | ABC zinc transporter permease ZnuB | -2.21 |
| *fepG* | PA14_10140 | ferric enterobactin transport protein FepG | -2.24 |
| *cioA* | PA14_13030 | CioA cyanide insensitive terminal oxidase | -2.30 |
| | PA14_71100 | hypothetical protein | -2.43 |
| | PA14_10920 | hypothetical protein | -2.68 |
| | PA14_02610 | phosphoribosyl dephospho CoA transferase | -2.71 |
| *dksA2* | PA14_73020 | DksA/TraR family C4 type zinc finger protein | -2.72 |
| *cntO* | PA14_63960 | CntO | -2.83 |
| *metE* | PA14_39590 | 5__methyltetrahydropteroyltriglutamate/homocysteine S methyltransferase | -3.05 |
| | | R-2L | |
| | PA14_64530 | hypothetical protein | 8.65 |
| | PA14_51050 | aldehyde dehydrogenase | 8.03 |
| *oprN* | PA14_32380 | multidrug efflux outer membrane protein OprN | 7.00 |
| | PA14_39060 | lipoprotein | 6.11 |
| | PA14_56660 | xenobiotic reductase | 5.21 |
| | PA14_51040 | oxidoreductase | 5.00 |
| | PA14_16660 | metal transporting Ptype ATPase | 4.41 |
| *mexF* | PA14_32390 | RND multidrug efflux transporter MexF | 4.30 |
| | PA14_22420 | hypothetical protein | 3.94 |
| | PA14_67540 | hypothetical protein | 3.39 |
| | PA14_28410 | hypothetical protein | 3.14 |
| | PA14_02740 | class II aldolase/adducin domain containing protein | 3.12 |
| | PA14_61180 | hypothetical protein | 2.90 |
| | PA14_71100 | hypothetical protein | 2.82 |

| | | | |
|---|---|---|---|
| | PA14_31950 | two component sensor | 2.45 |
| *pchF* | PA14_09280 | pyochelin synthetase | 2.44 |
| *arnF* | PA14_18310 | Probable 4-amino-4-deoxy-L-arabinose-phosphoundecaprenol flippase subunit | 2.38 |
| | PA14_73050 | GTP cyclohydrolase | 2.35 |
| | PA14_32640 | hypothetical protein | 2.26 |
| | PA14_18300 | nucleotide sugar dehydrogenase | 2.26 |
| *cntO* | PA14_63960 | CntO | 2.19 |
| *hslU* | PA14_66790 | ATP dependent protease ATP binding subunit HslU | 2.18 |
| *arnE* | PA14_18320 | Probable 4-amino-4-deoxy-L-arabinose-phosphoundecaprenol flippase subunit | 2.17 |
| | PA14_02930 | oxidoreductase | 2.14 |
| *arnT* | PA14_18330 | 4 amino 4 deoxy L arabinose transferase | 2.14 |
| *asrA* | PA14_54210 | ATP dependent protease | 2.13 |
| | PA14_47040 | TerC family protein | 2.13 |
| *glcC* | PA14_70710 | DNA binding transcriptional regulator GlcC | 2.12 |
| | PA14_54120 | ACP phosphodieterase | 2.08 |
| | PA14_58500 | hypothetical protein | 2.04 |
| *merD* | PA14_15450 | transcriptional regulator MerD | 2.02 |
| *opmE* | PA14_18790 | outer membrane efflux protein | 2.01 |
| | PA14_19370 | asparagine synthetase | -2.01 |
| *pilA* | PA14_58730 | type IV pilin structural subunit | -2.02 |
| *pcaK* | PA14_02900 | 4 hydroxybenzoate transporter PcaK | -2.03 |
| | PA14_26990 | hypothetical protein | -2.03 |
| *snr1* | PA14_24860 | cytochrome c Snr1 | -2.03 |
| | PA14_50280 | hypothetical protein | -2.04 |
| *PA14_21530* | PA14_21530 | ankyrin domain containing protein | -2.07 |
| *rhlG* | PA14_20270 | beta ketoacyl reductase | -2.07 |
| *oprM* | PA14_05550 | major intrinsic multiple antibiotic resistance efflux outer membrane protein OprM | -2.10 |
| | PA14_24960 | carbohydrate kinase | -2.14 |
| | PA14_36460 | hypothetical protein | -2.17 |

| | | | |
|---|---|---|---|
| | PA14_40540 | hypothetical protein | -2.18 |
| *cheW* | PA14_02230 | purine binding chemotaxis protein | -2.21 |
| | PA14_10270 | hypothetical protein | -2.25 |
| | PA14_19350 | hypothetical protein | -2.31 |
| *aer2* | PA14_02220 | chemotaxis transducer | -2.31 |
| | PA14_19360 | GNAT family acetyltransferase | -2.36 |
| *cheR2* | PA14_02200 | chemotaxis protein methyltransferase | -2.42 |
| *mexR* | PA14_05520 | multidrug resistance operon repressor MexR | -2.64 |
| *mexB* | PA14_05540 | RND multidrug efflux transporter MexB | -3.08 |
| | PA14_31090 | hypothetical protein | -3.26 |
| | PA14_02190 | hypothetical protein | -3.30 |
| *mexA* | PA14_05530 | RND multidrug efflux membrane fusion protein MexA | -4.00 |
| | | R-6E | |
| *armR* | PA14_16300 | mexR anti-repressor | 6.96 |
| | PA14_10900 | alcohol dehydrogenase | 6.47 |
| | PA14_10370 | hypothetical protein | 6.27 |
| | PA14_16290 | hypothetical protein | 6.22 |
| *actP* | PA14_22350 | acetate permease | 4.63 |
| | PA14_16310 | MFS permease | 4.06 |
| | PA14_33050 | hypothetical protein | 3.95 |
| | PA14_10400 | hypothetical protein | 3.78 |
| | PA14_22340 | hypothetical_protein | 3.72 |
| | PA14_33060 | hypothetical protein | 3.59 |
| *opdQ* | PA14_24790 | outer membrane porin | 3.58 |
| | PA14_37380 | flavin binding monooxygenase | 3.47 |
| | PA14_10380 | hypothetical protein | 3.33 |
| | PA14_18820 | hypothetical protein | 3.31 |
| *nalC* | PA14_16280 | transcriptional regulator | 3.25 |

| | | | |
|---|---|---|---|
| | PA14_23000 | ABC sugar transporter permease | 3.22 |
| *acsA* | PA14_52800 | acetyl CoA synthetase | 3.13 |
| | PA14_46610 | hypothetical protein | 3.12 |
| | PA14_55750 | chemotaxis transducer | 3.04 |
| | PA14_22990 | ABC sugar transporter permease | 2.98 |
| *gltK* | PA14_23010 | ABC transporter ATP binding protein | 2.92 |
| | PA14_47520 | transcriptional regulator | 2.90 |
| *nosR* | PA14_20230 | regulatory protein NosR | 2.89 |
| | PA14_47640 | major facilitator transporter | 2.86 |
| *pvdS* | PA14_33260 | extracytoplasmic function sigma 70 factor | 2.86 |
| *coxB* | PA14_01290 | cytochrome c oxidase subunit II | 2.82 |
| *norC* | PA14_06810 | nitric oxide reductase subunit C | 2.78 |
| | PA14_22380 | DNA polymerase III subunit epsilon | 2.74 |
| *pelG* | PA14_24560 | pelG hypothetical protein | 2.67 |
| *pelD* | PA14_24510 | pelD hypothetical protein | 2.58 |
| *pelE* | PA14_24530 | pelE hypothetical protein | 2.57 |
| | PA14_38610 | hypothetical protein | 2.53 |
| | PA14_06200 | hypothetical protein | 2.52 |
| *norB* | PA14_06830 | nitric oxide reductase subunit B | 2.46 |
| *oprB* | PA14_23030 | glucose/carbohydrate outer membrane porin OprB | 2.46 |
| | PA14_37360 | short chain dehydrogenase | 2.43 |
| *Eat* | PA14_11790 | amino acid transporter | 2.42 |
| *coxA* | PA14_01300 | cytochrome c oxidase subunit I | 2.41 |
| | PA14_49050 | hypothetical protein | 2.39 |
| | PA14_46620 | pyridine nucleotide disulfide oxidoreductase | 2.33 |
| | PA14_18830 | adenylosuccinate lyase | 2.32 |
| *pelF* | PA14_24550 | pelF hypothetical protein | 2.31 |
| *pelA* | PA14_24480 | pelA hypothetical protein | 2.31 |
| | PA14_18150 | acetyl coa synthetase | 2.29 |

| | | | |
|---|---|---|---|
| | PA14_20570 | chaperone | 2.28 |
| *bkdA1* | PA14_35530 | 2 oxoisovalerate dehydrogenase subunit alpha | 2.24 |
| | PA14_49690 | oxidoreductase | 2.23 |
| *gnyL* | PA14_38490 | hydroxymethylglutaryl Co lyase | 2.22 |
| | PA14_49310 | hypothetical protein | 2.22 |
| *bkdA2* | PA14_35520 | 2 oxoisovalerate dehydrogenase subunit beta | 2.22 |
| *fxsA* | PA14_57030 | FxsA protein | 2.22 |
| | PA14_39750 | amino acid permease | 2.21 |
| *opdP* | PA14_58410 | glycine glutamate dipeptide porin OpdP | 2.20 |
| *nosZ* | PA14_20200 | nitrous oxide reductase | 2.19 |
| *gnyH* | PA14_38470 | gamma carboxygeranoyl CoA hydratase | 2.17 |
| *pilX* | PA14_60300 | type 4 fimbrial biogenesis protein PilX | 2.16 |
| | PA14_11810 | aldehyde dehydrogenase | 2.16 |
| | PA14_18810 | hypothetical protein | 2.15 |
| | PA14_49300 | lipoxygenase | 2.10 |
| *gapA* | PA14_22890 | glyceraldehyde 3 phosphate dehydrogenase | 2.09 |
| *ibpA* | PA14_23680 | heat shock protein IbpA | 2.09 |
| | PA14_40540 | hypothetical protein | 2.09 |
| | PA14_02130 | hypothetical protein | 2.08 |
| | PA14_16410 | MFS transporter | 2.04 |
| | PA14_47610 | transcriptional regulator | 2.04 |
| *fimU* | PA14_60280 | type 4 fimbrial biogenesis protein FimU | 2.03 |
| | PA14_27490 | hypothetical protein | 2.02 |
| *bkdB* | PA14_35500 | branched chain alpha keto acid dehydrogenase subunit E2 | 2.02 |
| | PA14_46900 | hypothetical protein | 2.02 |
| *gnyA* | PA14_38480 | alpha subunit of geranoyl CoA carboxylase GnyA | 2.02 |
| | PA14_35330 | 2 ketogluconate transporter | 2.00 |
| *csaA* | PA14_22570 | CsaA protein | -2.00 |
| *phzF2* | PA14_39890 | phenazine biosynthesis protein | -2.02 |

| | | | |
|---|---|---|---|
| | PA14_28050 | chemotaxis transducer | -2.03 |
| | PA14_51950 | hypothetical protein | -2.04 |
| | PA14_39710 | radical SAM protein | -2.04 |
| *mexC* | PA14_60850 | multidrug efflux RND membrane fusion protein | -2.05 |
| | PA14_07990 | hypothetical protein | -2.06 |
| *pilA* | PA14_58730 | type IV pilin structural subunit | -2.06 |
| *lasA* | PA14_40290 | LasA protease | -2.07 |
| | PA14_08180 | hypothetical protein | -2.09 |
| *flgD* | PA14_50460 | flagellar basal body rod modification protein | -2.11 |
| *pa1L* | PA14_31290 | PA I galactophilic lectin | -2.11 |
| | PA14_37350 | hypothetical protein | -2.12 |
| *flgE* | PA14_50450 | flagellar hook protein FlgE | -2.12 |
| | PA14_61330 | magnesium transporter MgtC family | -2.12 |
| | PA14_20050 | outer membrane protein | -2.16 |
| | PA14_61540 | hypothetical protein | -2.16 |
| | PA14_46330 | transcriptional regulator | -2.16 |
| | PA14_63820 | hypothetical protein | -2.17 |
| | PA14_10910 | major facilitator transporter | -2.19 |
| | PA14_11320 | hypothetical protein | -2.19 |
| *motY* | PA14_18720 | OmpA family membrane protein | -2.19 |
| *cpbD* | PA14_53250 | chitin binding protein CbpD | -2.20 |
| *phnI* | PA14_20380 | hypothetical protein | -2.21 |
| | PA14_55820 | hypothetical protein | -2.22 |
| *hcp2* | PA14_43070 | Hcp2 | -2.22 |
| *aptA* | PA14_01620 | beta alanine pyruvate transaminase | -2.23 |
| *pctA* | PA14_56000 | chemotactic transducer PctA | -2.23 |
| *flgF* | PA14_50440 | flagellar basal body rod protein FlgF | -2.25 |
| | PA14_72960 | MFS dicarboxylate transporter | -2.26 |
| *opmD* | PA14_09500 | outer membrane protein | -2.26 |

| | | | |
|---|---|---|---|
| | PA14_40250 | outer membrane protein | -2.26 |
| *narG* | PA14_13780 | respiratory nitrate reductase alpha subun | -2.26 |
| | PA14_40520 | hypothetical protein | -2.27 |
| | PA14_34460 | hypothetical protein | -2.29 |
| | PA14_56670 | hypothetical protein | -2.30 |
| | PA14_61510 | hypothetical protein | -2.30 |
| | PA14_61500 | hypothetical protein | -2.32 |
| *pprA* | PA14_55780 | two component sensor | -2.32 |
| | PA14_36560 | hypothetical protein | -2.32 |
| | PA14_22880 | Fe S protein | -2.34 |
| *prpL* | PA14_09900 | Pvds regulated endoprotease lysyl class | -2.35 |
| | PA14_29800 | chemotaxis transducer | -2.35 |
| *phzD1* | PA14_09450 | phenazine biosynthesis protein PhzD | -2.35 |
| | PA14_61530 | pili assembly chaperone | -2.35 |
| *fliK* | PA14_45830 | hypothetical protein | -2.37 |
| *msuE* | PA14_34180 | NADH dependent FMN reductase MsuE | -2.39 |
| *flgC* | PA14_50470 | flagellar basal body rod protein FlgC | -2.42 |
| *pncA* | PA14_64950 | hypothetical protein | -2.45 |
| *ureF* | PA14_64660 | urease accessory protein UreF | -2.47 |
| *rcpA* | PA14_55920 | type II secretion system protein | -2.48 |
| | PA14_40240 | ABC transporter ATP binding protein/permease | -2.50 |
| *pvdG* | PA14_33270 | protein PvdG | -2.51 |
| | PA14_55860 | hypothetical protein | -2.51 |
| *tagT1* | PA14_00860 | TagT1 | -2.51 |
| *hxcX* | PA14_55480 | HxcX | -2.55 |
| *dkgB* | PA14_09980 | 2,5 diketo D gluconate reductase B | -2.58 |
| *phnK* | PA14_20400 | phosphonate C P lyase system protein PhnK | -2.59 |
| | PA14_71100 | hypothetical protein | -2.64 |
| | PA14_61520 | hypothetical protein | -2.69 |

| | | | |
|---|---|---|---|
| *narK1* | PA14_13750 | nitrite extrusion protein1 | -2.69 |
| | PA14_55900 | hypothetical protein | -2.70 |
| *narK2* | PA14_13770 | nitrite extrusion protein2 | -2.70 |
| *phnJ* | PA14_20390 | hypothetical protein | -2.72 |
| *cntO* | PA14_63960 | CntO | -2.73 |
| | PA14_55890 | type II secretion system protein | -2.77 |
| *lasI* | PA14_45940 | autoinducer synthesis protein LasI | -2.81 |
| | PA14_55850 | pilus assembly protein | -2.81 |
| *phnG* | PA14_20360 | phosphonate metabolism protein PhnG | -2.82 |
| | PA14_55880 | hypothetical protein | -2.84 |
| | PA14_34170 | hypothetical protein | -2.85 |
| | PA14_40260 | hypothetical protein | -2.87 |
| *phoA* | PA14_21410 | alkaline phosphatase | -2.89 |
| | PA14_55930 | pilus assembly protein | -2.89 |
| | PA14_39090 | hypothetical protein | -2.93 |
| *phnH* | PA14_20370 | carbon phosphorus lyase complex subunit | -2.95 |
| | PA14_55840 | hypothetical protein | -3.04 |
| | PA14_73020 | DksA/TraR family C4 type zinc finger protein | -3.04 |
| | PA14_66510 | MFS transporter | -3.05 |
| | PA14_30890 | hypothetical protein | -3.12 |
| *metE* | PA14_39590 | 5 methyltetrahydropteroyltriglutamate/homocysteine S methyltransferase | -3.16 |
| | PA14_55940 | hypothetical protein | -3.26 |
| *hasAp* | PA14_20020 | heme acquisition protein HasAp | -3.29 |
| *moeA1* | PA14_13280 | molybdenum cofactor biosynthetic protein A1 | -3.51 |
| *moaB1* | PA14_13260 | MoaB1 | -3.57 |
| | PA14_63770 | hypothetical protein | -4.21 |
| *phoU* | PA14_70800 | phosphate uptake regulatory protein PhoU | -4.43 |
| *pstA* | PA14_70830 | phosphate ABC transporter permease | -4.46 |
| *pstC* | PA14_70850 | membrane protein component of ABC phosphate transporter | -4.52 |

| | | | |
|---|---|---|---|
| | PA14_71110 | lipolytic protein | -4.73 |
| | PA14_39780 | hypothetical protein | -4.75 |
| *mgtA* | PA14_63800 | Mg(2+) transport ATPase P type 2 | -4.81 |
| | PA14_63750 | hypothetical protein | -5.03 |
| | PA14_19680 | hypothetical protein | -5.20 |
| *pstB* | PA14_70810 | phosphate transporter ATP binding protein | -5.34 |
| | PA14_19690 | hypothetical protein | -5.34 |
| | PA14_70860 | hypothetical protein | -5.42 |
| | PA14_63780 | hypothetical protein | -5.71 |
| | PA14_63740 | hypothetical protein | -6.05 |
| | | **R-6L** | |
| | PA14_16290 | hypothetical protein | 13.98 |
| *armR* | PA14_16300 | mexR anti-repressor | 13.70 |
| | PA14_16310 | MFS permease | 12.44 |
| *pchF* | PA14_09280 | pyochelin synthetase | 9.73 |
| *pchG* | PA14_09290 | pyochelin biosynthetic protein PchG | 7.98 |
| *pchE* | PA14_09270 | dihydroaeruginoic acid synthetase | 7.28 |
| | PA14_09380 | transporter | 7.13 |
| *pchB* | PA14_09220 | isochorismate pyruvate lyase | 6.82 |
| *pchD* | PA14_09240 | pyochelin biosynthesis protein PchD | 6.66 |
| *pchA* | PA14_09210 | salicylate biosynthesis isochorismate synthase | 6.63 |
| | PA14_09300 | ABC transporter ATP binding protein | 6.44 |
| *fptA* | PA14_09340 | Fe(III) pyochelin outer membrane receptor | 6.41 |
| *pchC* | PA14_09230 | pyochelin biosynthetic protein PchC | 6.23 |
| | PA14_09350 | hypothetical protein | 6.07 |
| | PA14_16280 | transcriptional regulator | 5.32 |
| | PA14_51050 | aldehyde dehydrogenase | 4.93 |
| | PA14_09370 | hypothetical protein | 4.56 |
| | PA14_09320 | ABC transporter ATP binding protein | 4.04 |

| | | | |
|---|---|---|---|
| | PA14_10370 | hypothetical protein | 3.62 |
| | PA14_16660 | metal transporting P type ATPase | 3.61 |
| | PA14_51040 | oxidoreductase | 3.59 |
| | PA14_58500 | hypothetical protein | 3.49 |
| *antC* | PA14_32140 | anthranilate dioxygenase reductase | 3.34 |
| | PA14_71100 | hypothetical protein | 3.27 |
| *czcS* | PA14_31950 | two component sensor | 3.24 |
| *antB* | PA14_32150 | anthranilate dioxygenase small subunit | 3.14 |
| *antA* | PA14_32160 | anthranilate dioxygenase large subunit | 3.00 |
| | PA14_14560 | hypothetical protein | 2.88 |
| | PA14_47520 | transcriptional regulator | 2.86 |
| | PA14_72360 | hypothetical protein | 2.79 |
| | PA14_23000 | ABC sugar transporter permease | 2.77 |
| | PA14_18300 | nucleotide sugar dehydrogenase | 2.77 |
| | PA14_37630 | amino acid permease | 2.77 |
| | PA14_61200 | hypothetical protein | 2.77 |
| *gltK* | PA14_23010 | ABC transporter ATP binding protein | 2.75 |
| | PA14_37610 | kynureninase | 2.72 |
| | PA14_38950 | hypothetical protein | 2.72 |
| | PA14_36920 | hypothetical protein | 2.71 |
| | PA14_07470 | tRNA Met | 2.69 |
| *betT1* | PA14_70980 | choline transporter BetT | 2.60 |
| *pelG* | PA14_24560 | hypothetical protein | 2.56 |
| *pvdN* | PA14_33720 | protein PvdN | 2.56 |
| *mreD* | PA14_58120 | rod shape determining protein MreD | 2.47 |
| | PA14_22990 | ABC sugar transporter permease | 2.46 |
| | PA14_22980 | sugar ABC transporter substrate binding protein | 2.45 |
| | PA14_67140 | hypothetical protein | 2.43 |
| *pelB* | PA14_24490 | hypothetical protein | 2.42 |

| | | | |
|---|---|---|---|
| *dguA* | PA14_67150 | oxidoreductase | 2.40 |
| *dguC* | PA14_67130 | ABC transporter substrate binding protein | 2.40 |
| *pelA* | PA14_24480 | hypothetical protein | 2.39 |
| | PA14_20480 | hypothetical protein | 2.37 |
| *pvdQ* | PA14_33820 | penicillin acylase related protein | 2.36 |
| | PA14_72370 | hypothetical protein | 2.35 |
| *ligD* | PA14_36910 | ATP dependent DNA ligase | 2.35 |
| *mexA* | PA14_05530 | RND multidrug efflux membrane fusion protein MexA | 2.30 |
| *cdrB* | PA14_61190 | hypothetical protein | 2.26 |
| | PA14_72710 | transporter | 2.26 |
| | PA14_20470 | hypothetical protein | 2.23 |
| *pvdO* | PA14_33710 | protein PvdO | 2.23 |
| | PA14_72700 | hypothetical protein | 2.23 |
| *mexB* | PA14_05540 | RND multidrug efflux transporter MexB | 2.22 |
| *phzA1* | PA14_09480 | phenazine biosynthesis protein | 2.22 |
| *mqoB* | PA14_61400 | malate:quinone oxidoreductase | 2.19 |
| | PA14_55050 | TonB dependent receptor | 2.18 |
| *pelD* | PA14_24510 | hypothetical protein | 2.18 |
| | PA14_20460 | hypothetical protein | 2.17 |
| | PA14_64430 | hypothetical protein | 2.14 |
| | PA14_55040 | ferric enterobactin transporter ATP binding protein | 2.11 |
| | PA14_04180 | hypothetical protein | 2.11 |
| | PA14_37780 | hypothetical protein | 2.11 |
| *kynB* | PA14_37590 | kynurenine formamidase KynB | 2.10 |
| | PA14_47040 | TerC family protein | 2.10 |
| | PA14_28600 | hypothetical protein | 2.10 |
| *hisC2* | PA14_23290 | histidinol phosphate aminotransferase | 2.10 |
| *oprB* | PA14_23030 | glucose/carbohydrate outer membrane porin OprB | 2.09 |
| *pdxA* | PA14_07740 | 4 hydroxythreonine 4 phosphate dehydrogenase | 2.09 |

| | | | |
|---|---|---|---|
| | PA14_37760 | MFS transporter | 2.09 |
| | PA14_33050 | hypothetical protein | 2.06 |
| *tse3* | PA14_19020 | Tse3 | 2.06 |
| | PA14_15750 | hypothetical protein | 2.05 |
| | PA14_46640 | siderophore receptor | 2.05 |
| | PA14_54040 | amino acid permease | 2.03 |
| *betA* | PA14_70940 | choline dehydrogenase | 2.03 |
| *pelE* | PA14_24530 | hypothetical protein | 2.02 |
| | PA14_56170 | hypothetical protein | 2.02 |
| | PA14_38130 | amino acid permease | 2.01 |
| *czcR* | PA14_31960 | two component response regulator | 2.01 |
| | PA14_29300 | transcriptional regulator | -2.00 |
| | PA14_39710 | radical SAM protein | -2.00 |
| | PA14_20840 | hypothetical protein | -2.00 |
| | PA14_13840 | peptidyl prolyl cis trans isomerase PpiC type | -2.00 |
| | PA14_30500 | acyl CoA dehydrogenase | -2.01 |
| *ssuB* | PA14_19580 | aliphatic sulfonates transport ATP binding subunit | -2.01 |
| | PA14_24950 | FAD dependent glycerol 3 phosphate dehydrogenase | -2.01 |
| | PA14_02190 | hypothetical protein | -2.01 |
| | PA14_64530 | hypothetical protein | -2.02 |
| | PA14_52910 | hypothetical protein | -2.02 |
| | PA14_19530 | NAD(P)H dependent FMN reductase | -2.03 |
| | PA14_37470 | flavin dependent oxidoreductase | -2.04 |
| | PA14_61950 | hypothetical protein | -2.04 |
| | PA14_12450 | acyl CoA dehydrogenase | -2.04 |
| *pemB* | PA14_44480 | PemB | -2.05 |
| *cgrA* | PA14_37070 | hypothetical protein | -2.05 |
| *pctA* | PA14_56000 | chemotactic transducer PctA | -2.05 |
| *morB* | PA14_26130 | morphinone reductase | -2.05 |

| | | | |
|---|---|---|---|
| *pncA* | PA14_64950 | hypothetical protein | -2.07 |
| | PA14_26110 | MFS transporter | -2.07 |
| | PA14_71740 | pyruvate carboxylase subunit A | -2.08 |
| | PA14_51830 | DNA binding stress protein | -2.08 |
| *phzF1* | PA14_09420 | phenazine biosynthesis protein | -2.08 |
| | PA14_52110 | hypothetical protein | -2.09 |
| | PA14_39670 | hypothetical protein | -2.09 |
| | PA14_71720 | pyruvate carboxylase subunit B | -2.09 |
| | PA14_19920 | branched chain alpha keto acid dehydrogenase subunit E2 | -2.09 |
| *fliC* | PA14_50290 | flagellin type B | -2.10 |
| | PA14_20980 | short chain dehydrogenas | -2.11 |
| *adhA* | PA14_71630 | alcohol dehydrogenase | -2.12 |
| | PA14_21010 | FAD dependent monooxygenase | -2.13 |
| | PA14_29320 | NADH dehydrogenase FAD containing subunit | -2.14 |
| *nirQ* | PA14_06770 | regulatory protein NirQ | -2.15 |
| | PA14_12470 | hypothetical protein | -2.15 |
| *narK2* | PA14_13770 | nitrite extrusion protein 2 | -2.15 |
| | PA14_32350 | hypothetical protein | -2.16 |
| | PA14_02460 | NAD(P) transhydrogenase subunit alpha part 2 | -2.17 |
| | PA14_58720 | hypothetical protein | -2.17 |
| | PA14_55930 | pilus assembly protein | -2.18 |
| | PA14_21260 | hypothetical protein | -2.18 |
| | PA14_02490 | tonB2 hypothetical protein | -2.20 |
| | PA14_72260 | hypothetical protein | -2.20 |
| | PA14_12920 | taurine ABC transporter periplasmic protein | -2.21 |
| | PA14_20280 | hypothetical protein | -2.22 |
| | PA14_21540 | 3 oxoacyl ACP synthase | -2.22 |
| | PA14_48830 | transcriptional regulator | -2.23 |
| | PA14_19540 | hypothetical protein | -2.23 |

| | | | |
|---|---|---|---|
| | PA14_02450 | NAD(P) transhydrogenase subunit alpha part 1 | -2.23 |
| | PA14_24960 | carbohydrate kinase | -2.23 |
| | PA14_40270 | cation transporter | -2.23 |
| | PA14_55790 | hypothetical protein | -2.24 |
| | PA14_66460 | hypothetical protein | -2.24 |
| *rfaD* | PA14_20890 | ADP L glycero D manno heptose 6 epimerase | -2.24 |
| | PA14_32660 | MFS family transporter | -2.24 |
| | PA14_58690 | hypothetical protein | -2.26 |
| *opmD* | PA14_09500 | outer membrane protein | -2.29 |
| | PA14_48510 | hypothetical protein | -2.29 |
| *mtlY* | PA14_34350 | xylulose kinase | -2.30 |
| *lasA* | PA14_40290 | LasA protease | -2.30 |
| | PA14_63750 | hypothetical protein | -2.32 |
| | PA14_46800 | hypothetical protein | -2.33 |
| | PA14_48140 | hypothetical protein | -2.35 |
| | PA14_60560 | hypothetical protein | -2.35 |
| | PA14_64620 | oxidoreductase | -2.36 |
| | PA14_51950 | hypothetical protein | -2.37 |
| | PA14_61010 | hypothetical protein | -2.38 |
| *pilA* | PA14_58730 | type IV pilin structural subunit | -2.39 |
| *narG* | PA14_13780 | respiratory nitrate reductase alpha subunit | -2.42 |
| | PA14_10910 | major facilitator transporter | -2.43 |
| | PA14_50280 | hypothetical protein | -2.46 |
| *katB* | PA14_61040 | catalase | -2.51 |
| | PA14_05860 | hypothetical protein | -2.51 |
| *lldP* | PA14_63080 | L lactate permease | -2.53 |
| *snr1* | PA14_24860 | cytochrome c Snr1 | -2.55 |
| | PA14_02200 | chemotaxis protein methyltransferase | -2.58 |
| | PA14_61000 | hypothetical protein | -2.66 |

| | | | |
|---|---|---|---|
| | PA14_05880 | membrane bound protease | -2.70 |
| *nirM* | PA14_06740 | cytochrome c 551 | -2.73 |
| | PA14_39420 | hypothetical protein | -2.73 |
| *cheA* | PA14_02250 | two component sensor | -2.77 |
| | PA14_02220 | chemotaxis transducer | -2.78 |
| *cheB* | PA14_02260 | two component response regulator | -2.88 |
| *bfrA* | PA14_09160 | bacterioferritin | -2.88 |
| *phzG2* | PA14_39880 | pyridoxamine 5'phosphate oxidase | -2.89 |
| | PA14_30890 | hypothetical protein | -2.90 |
| | PA14_53300 | alkyl hydroperoxide reductase | -2.92 |
| | PA14_26990 | hypothetical protein | -2.96 |
| *ssuD* | PA14_19560 | alkanesulfonate monooxygenase | -2.99 |
| *cheW* | PA14_02230 | purine binding chemotaxis protein | -3.00 |
| | PA14_63780 | hypothetical protein | -3.00 |
| *nirE* | PA14_06660 | uroporphyrin III c methyltransferase | -3.03 |
| | PA14_27290 | hypothetical protein | -3.03 |
| | PA14_63740 | hypothetical protein | -3.04 |
| *metE* | PA14_39590 | 5 methyltetrahydropteroyltriglutamate/homocysteine S methyltransferase | -3.15 |
| | PA14_63770 | hypothetical protein | -3.15 |
| *ahpF* | PA14_01720 | alkyl hydroperoxide reductase | -3.26 |
| | PA14_55940 | hypothetical protein | -3.32 |
| *nirJ* | PA14_06670 | heme d1 biosynthesis protein NirJ | -3.36 |
| *trxB2* | PA14_53290 | thioredoxin reductase 2 | -3.55 |
| *ureE* | PA14_64650 | urease accessory protein UreE | -3.58 |
| | PA14_21530 | ankyrin domain containing protein | -3.71 |
| | PA14_03090 | hypothetical protein | -3.74 |
| | PA14_39730 | hypothetical protein | -3.75 |
| *katA* | PA14_09150 | catalase | -3.86 |
| *nirH* | PA14_06680 | hypothetical protein | -4.00 |

| | | | |
|---|---|---|---|
| | PA14_03080 | acetyltransferase | -4.31 |
| *nosL* | PA14_20150 | NosL protein | -4.35 |
| *nirG* | PA14_06690 | transcriptional regulator | -4.40 |
| | PA14_61020 | hypothetical protein | -4.48 |
| *nirF* | PA14_06720 | heme d1 biosynthesis protein NirF | -4.73 |
| *nirL* | PA14_06700 | heme d1 biosynthesis protein NirL | -4.74 |
| | PA14_22880 | Fe S protein | -4.78 |
| | PA14_42000 | hypothetical protein | -4.86 |
| | PA14_06790 | cytochrome c oxidase subunit | -5.10 |
| | PA14_22320 | hypothetical protein | -5.22 |
| *nosY* | PA14_20170 | NosY protein | -5.59 |
| | PA14_06860 | hypothetical protein | -5.59 |
| | PA14_06800 | hypothetical protein | -5.77 |
| | PA14_21090 | hypothetical protein | -5.88 |
| *nosD* | PA14_20190 | copper ABC transporter periplasmic substrate binding protein | -6.21 |
| | PA14_06710 | transcriptional regulator | -6.34 |
| *nosR* | PA14_20230 | regulatory protein NosR | -7.05 |
| *nirC* | PA14_06730 | c type cytochrome | -7.53 |
| | PA14_55840 | hypothetical protein | -7.54 |
| *nosZ* | PA14_20200 | nitrous oxide reductase | -9.12 |
| *nosF* | PA14_20180 | NosF protein | -9.74 |
| *norC* | PA14_06810 | nitric oxide reductase subunit C | -11.89 |
| *norB* | PA14_06830 | nitric oxide reductase subunit B | -13.52 |
| *norD* | PA14_06840 | dinitrification protein NorD | -14.60 |

**S2 Table: Plasmids Used in This Study**

| Plasmid Name | Description | Primers Used |
|---|---|---|
| pMini-CTX-lacZ | promoterless lacZ, integration plasmid for PA14 | - |
| pFLP2 | flip recombinase for Mini-CTX clean up | - |
| pEXG2tc | PA14 deletion plasmid, empty vector | - |
| pKN03 | Mini-CTX-$P_{armR}$-lacZ | ECP031/ECP032 |
| pKN08 | EXG2tc Δ*mexAB-oprM* | ECP047- ECP050 |
| pKN09 | EXG2tc Δ*mexEF-oprN* | ECP051-ECP054 |
| pHL03 | EXG2tc Δ*mexCD-oprJ* | ECP091-ECP094 |
| pHL04 | EXG2tc Δ*mexXY* | ECP095-ECP098 |
| pHL05 | Mini-CTX-$P_{mexAB\text{-}oprM}$-lacZ | ECP120-ECP121 |

**S3 Table: Cloning Primers Used in This Study**

| Primer Name | Sequence (5' → 3') | Description |
|---|---|---|
| ECP031 | ATCGCTAGTTAGTTAGGATCGGGGGACTCCTGCGG GGAGG | Mini-CTX-$P_{armR}$-lacZ FOR |
| ECP032 | GATAAGCTTGATATCGAATTGGCGGTATCGGGCCTC GCGG | Mini-CTX-$P_{armR}$-lacZ REV |
| ECP047 | GCATAAATGTAAAGCAAGCTCGACGAGAAGAACCC GTCGG | Clone US region of *mexAB-oprM* into pEXG2tc FOR |
| ECP048 | TTGCATGGCGCGGAAGGCGAATTCAGAACCTGAAA CAAGGTTGA | Clone US region of *mexAB-oprM* into pEXG2tc REV |
| ECP049 | CCTTGTTTCAGGTTCTGAATTCGCCTTCCGCGCCAT GCAAG | Clone DS region of *mexAB-oprM* into pEXG2tc FOR |
| ECP050 | ATTAATTAAGGTACCGAATTGTACCGGCCGCGCCG GACGC | Clone DS region of *mexAB-oprM* into pEXG2tc REV |
| ECP051 | GCATAAATGTAAAGCAAGCTCGCTGTTCGACGACC CGCTG | Clone US region of *mexEF-oprN* into pEXG2tc FOR |
| ECP052 | ATCGGCCGGGGATAGCCGGTGCTTGACTCCGCCAG TCGGT | Clone US region of *mexEF-oprN* into pEXG2tc REV |
| ECP053 | ACCGACTGGCGGAGTCAAGCACCGGCTATCCCCGG CCGAT | Clone DS region of *mexEF-oprN* into pEXG2tc FOR |
| ECP054 | ATTAATTAAGGTACCGAATTGCGTCGCTGGCGCATC GGCT | Clone DS region of *mexEF-oprN* into pEXG2tc REV |
| ECP091 | GCATAAATGTAAAGCAAGCTGGAGGTGTGATTCGG TAGAC | Clone US region of *mexCD-oprJ* into pEXG2tc FOR |
| ECP092 | TTTCCCACACGTTTACCCGCGACACACCCGACCGTT GATT | Clone US region of *mexCD-oprJ* into pEXG2tc REV |
| ECP093 | AATCAACGGTCGGGTGTGTCGCGGGTAAACGTGTG GGAAA | Clone DS region of *mexCD-oprJ* into pEXG2tc FOR |
| ECP094 | ATTAATTAAGGTACCGAATTTGTTCGCCGTCCGATT CCAG | Clone DS region of *mexCD-oprJ* into pEXG2tc REV |
| ECP095 | GCATAAATGTAAAGCAAGCTGCAGCCAAGCGCGGT GGCAG | Clone US region of *mexXY* into pEXG2tc FOR |
| ECP096 | CGAGCGCTGCCGGCGGCCGAGGGTGTCCCTCGATT CGTGAACTCG | Clone US region of *mexXY* into pEXG2tc REV |
| ECP097 | TCACGAATCGAGGGACACCCTCGGCCGCCGGCAGC GCTCG | Clone DS region of *mexXY* into pEXG2tc FOR |
| ECP098 | GGTACCGAATTGACAGCCGTGAAAGATACTTGTCA GAAATATTTAATCGGCGCC | Clone DS region of *mexXY* into pEXG2tc REV |
| ECP120 | ATCGCTAGTTAGTTAGGATCATTCAGAACCTGAAAC AAGGT | Clone pMini-CTX-$P_{mexAB\text{-}oprM}$-lacZ FOR |
| ECP121 | GATAAGCTTGATATCGAATTTGGTTTGGCCGAGTAA ACCT | Clone pMini-CTX-$P_{mexAB\text{-}oprM}$-lacZ REV |

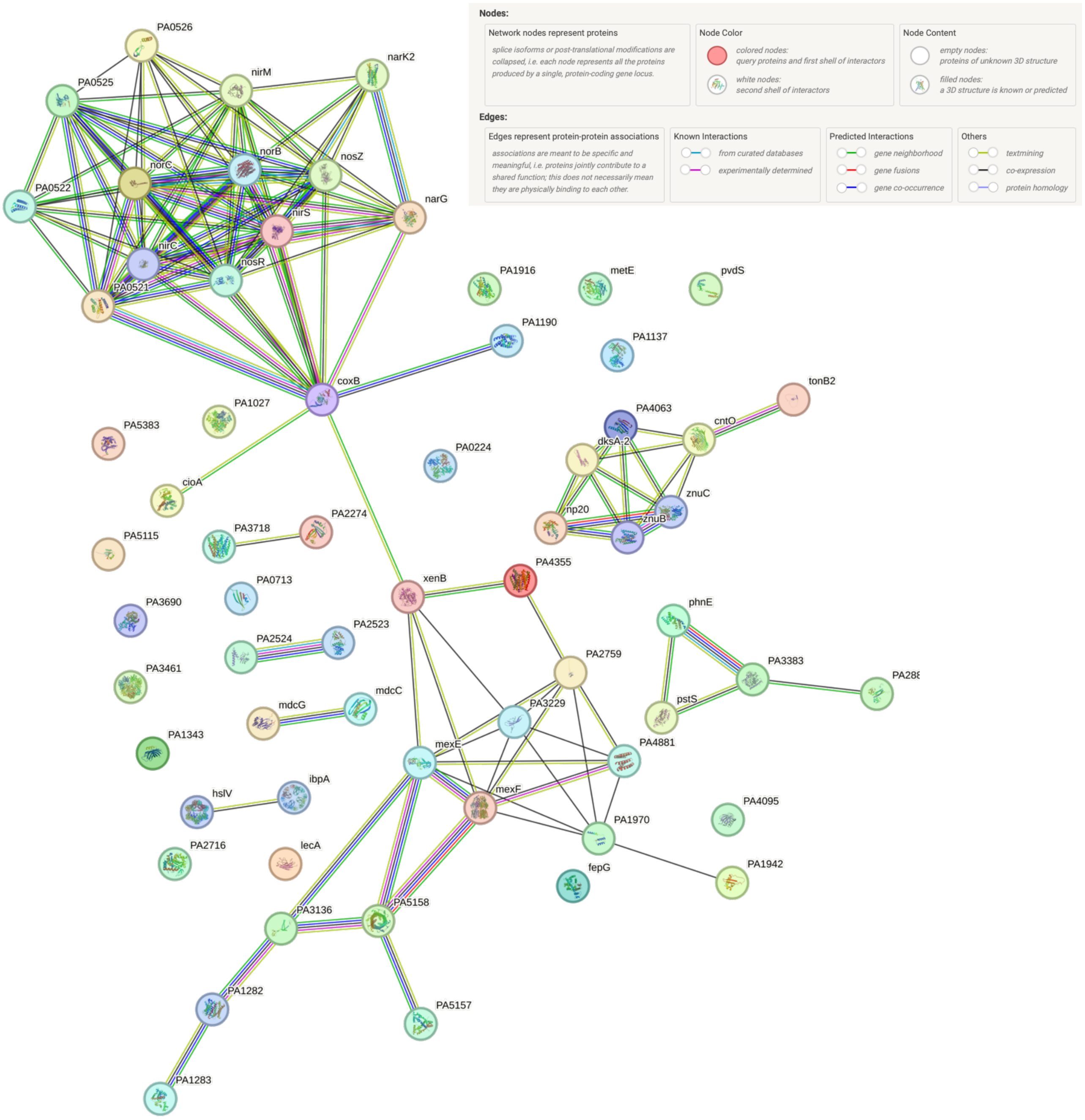


**S1 Fig: R-2E STRING Analysis.** Figure created in STRING Database (STRING Consortium 2024, www.string-db.org) from differentially expressed genes in R-2E treatment compared to DMSO. PPI enrichment p-value <1.0e-16, reference genome PAO1.

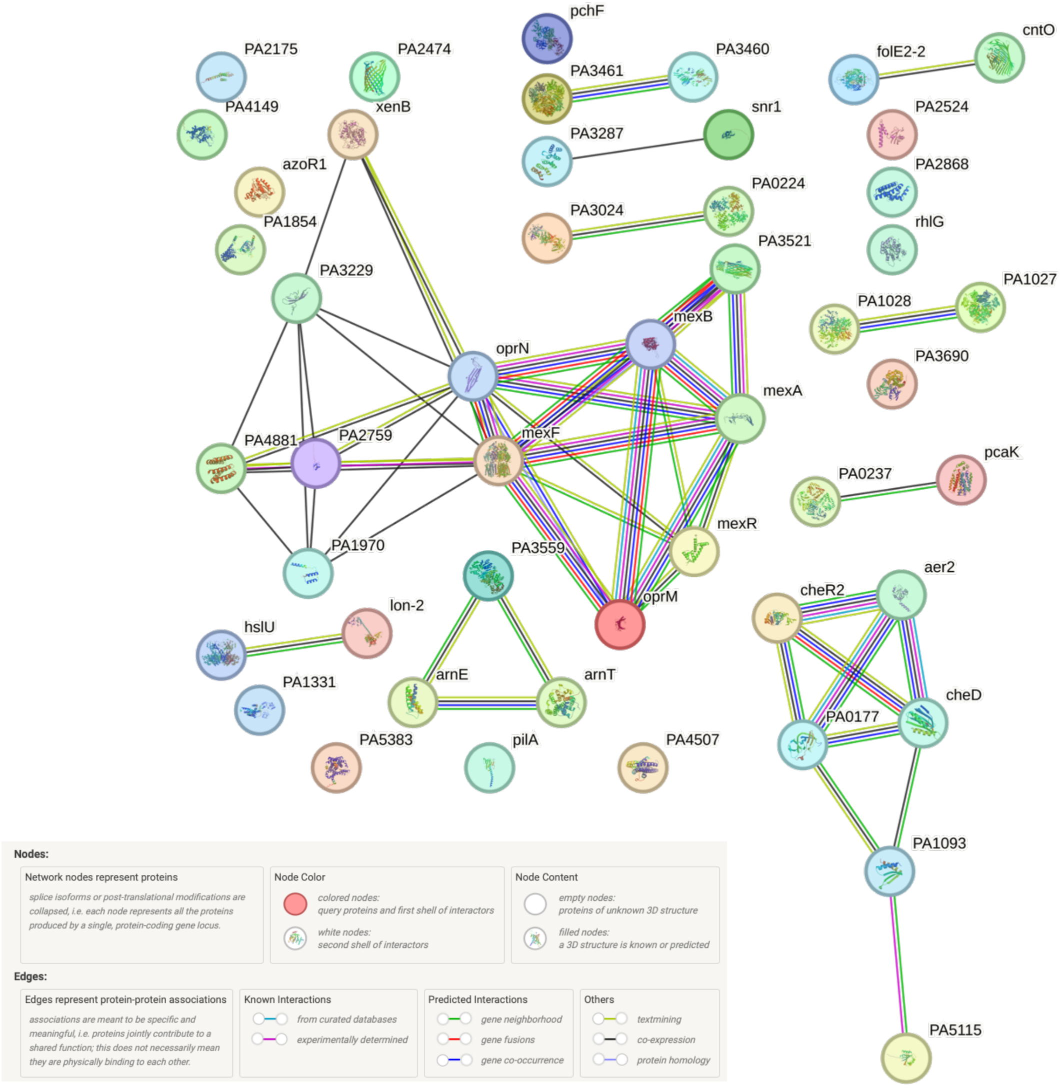


**S2 Fig: R-2L STRING Analysis.** Figure created in STRING Database (STRING Consortium 2024, www.string-db.org) from differentially expressed genes in R-2L treatment compared to DMSO. PPI enrichment p-value <1.0e-16, reference genome PAO1.

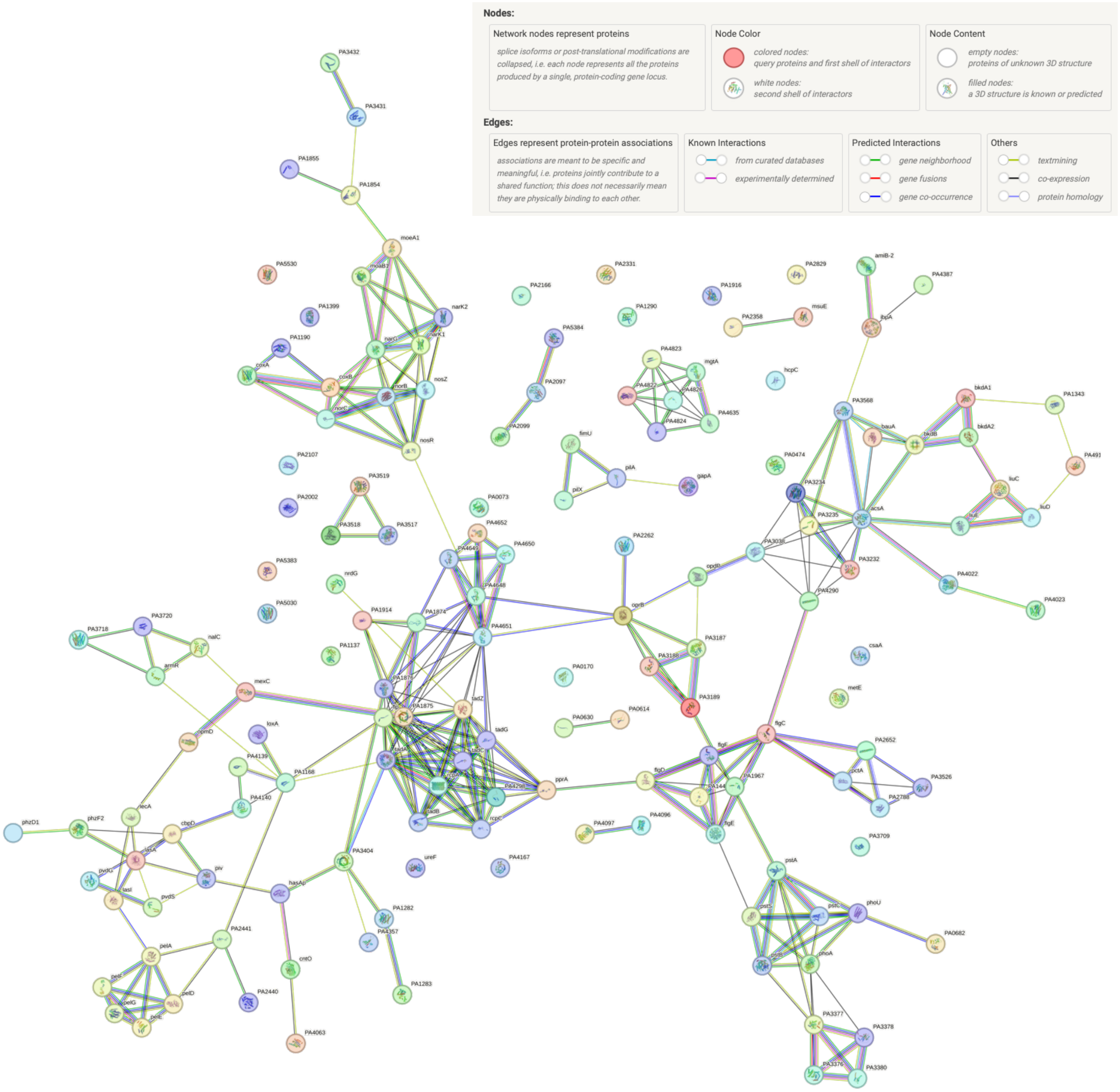


**S3 Fig: R-6E STRING Analysis.** Figure created in STRING Database (STRING Consortium 2024, www.string-db.org) from differentially expressed genes in R-6E treatment compared to DMSO. PPI enrichment p-value <1.0e-16, reference genome PAO1.

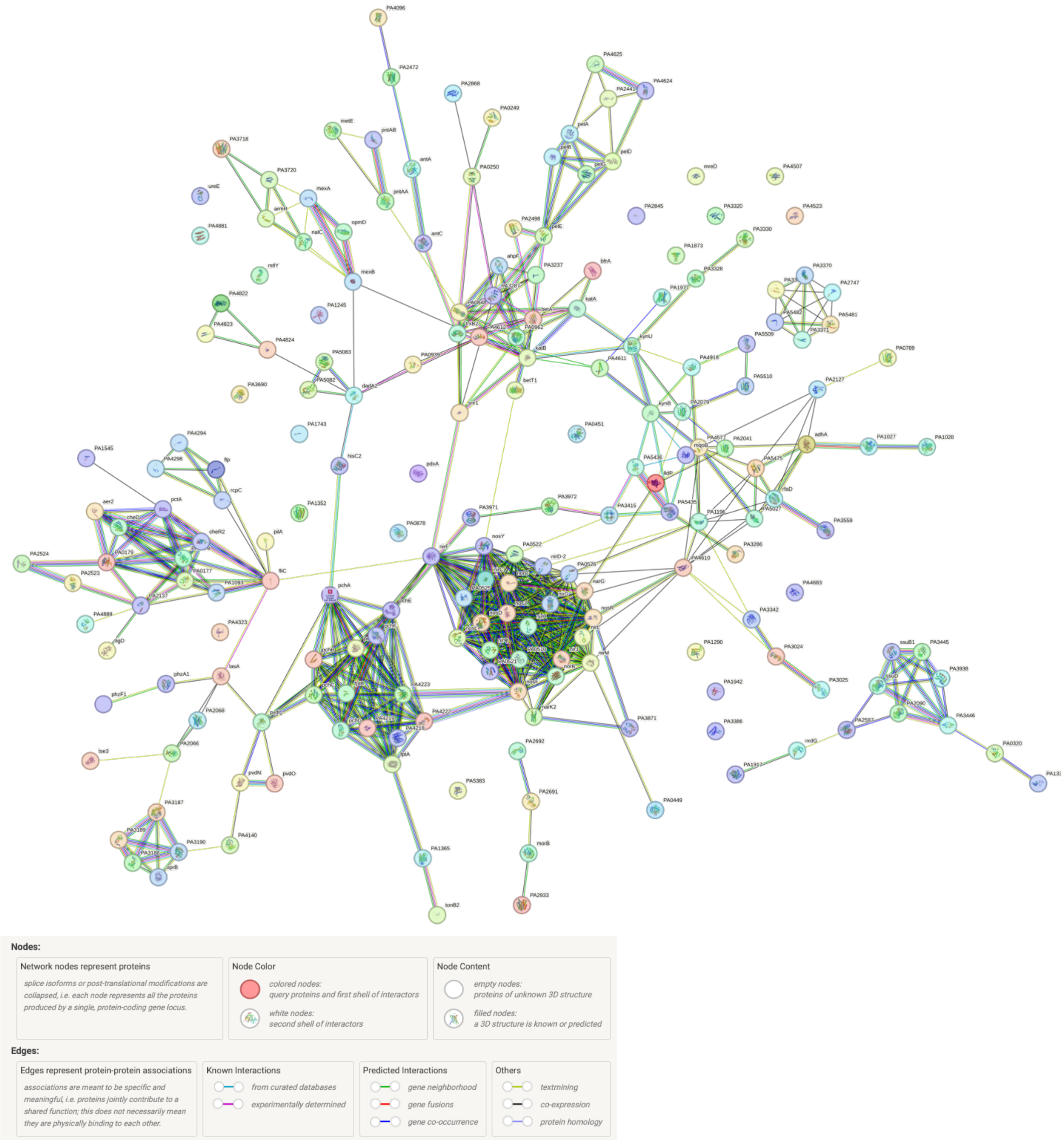


**S4 Fig: R-6L STRING Analysis.** Figure created in STRING Database (STRING Consortium 2024, www.string-db.org) from differentially expressed genes in R-6L treatment compared to DMSO. PPI enrichment p-value <1.0e-16, reference genome PAO1.

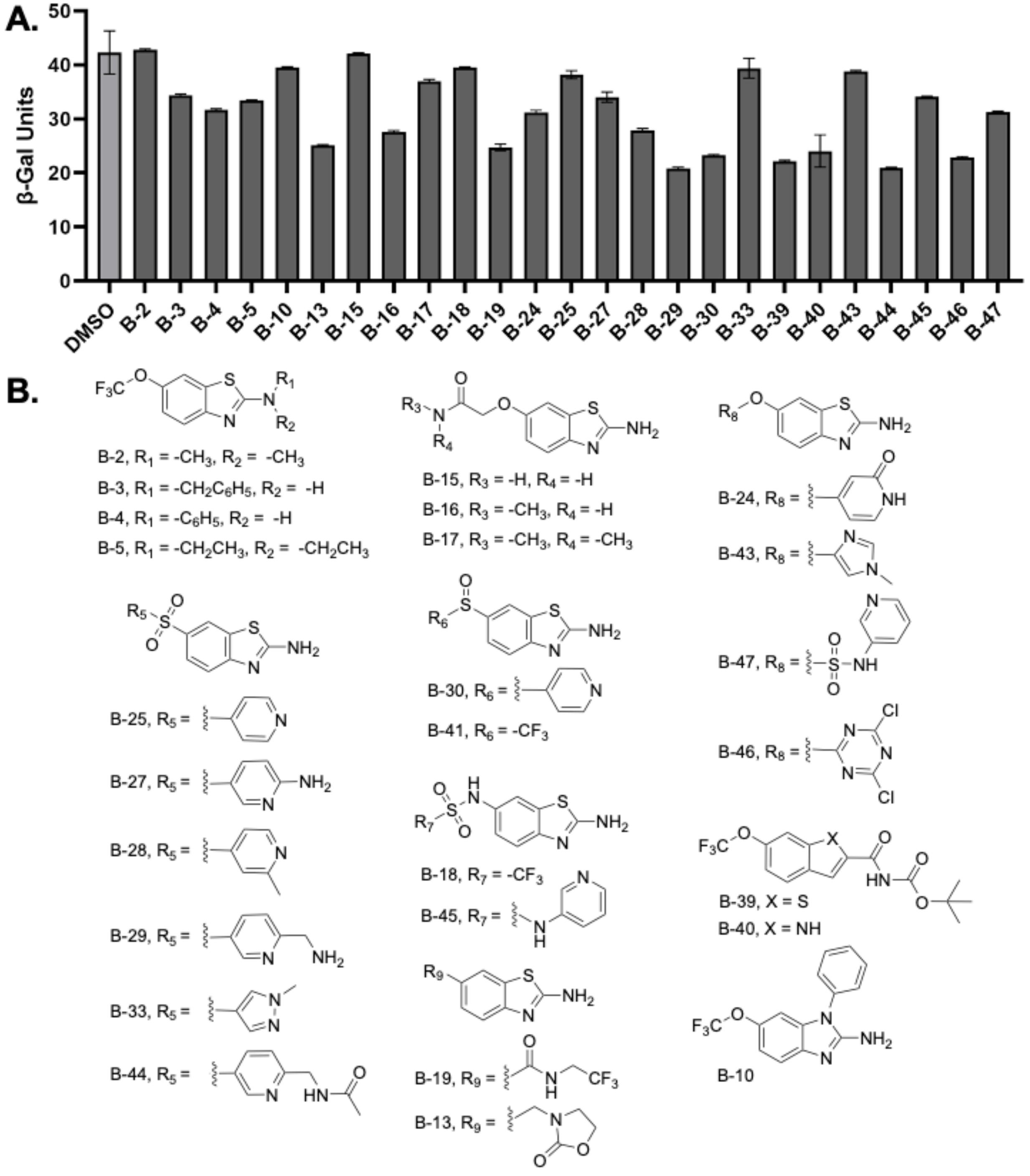


**S5 Fig: B-Series Non-Hit Compound Structures and β-Gal Results.** A. β-galactosidase units for each B-series inhibitor that did not meet the threshold of 20 units above or below the DMSO value in the PA14::$P_{armR}$-lacZ strain. All inhibitors were treated at 200 µM. B. Structures of the B-series inhibitors that were designated "non-hits".

**S4 Table: MIC Values**

| Strain | MIC µg/mL (µM) | | | |
|---|---|---|---|---|
| | **R-2*** | **R-6** | **Imipenem** | **Ciprofloxacin** |
| WT | >630 (2000) | 700 (3200) | 1 | 0.25 |
| ΔAB | >630 (2000) | 700 (3200) | - | 0.06 |
| ΔCD | >630 (2000) | 700 (3200) | - | 0.13 |
| ΔEF | >630 (2000) | 700 (3200) | - | 0.13 |
| ΔXY | >630 (2000) | 700 (3200) | - | 0.25 |
| $\Delta^2$ | >630 (2000) | 350(1600) | - | 0.06 |
| $\Delta^3$ | >630 (2000) | 87 (400) | - | 0.03 |
| $\Delta^4$ | >630 (2000) | 87 (400) | - | <0.004 |

*2000 µM (630 µg/mL) is the solubility limit for R-2 in LB media

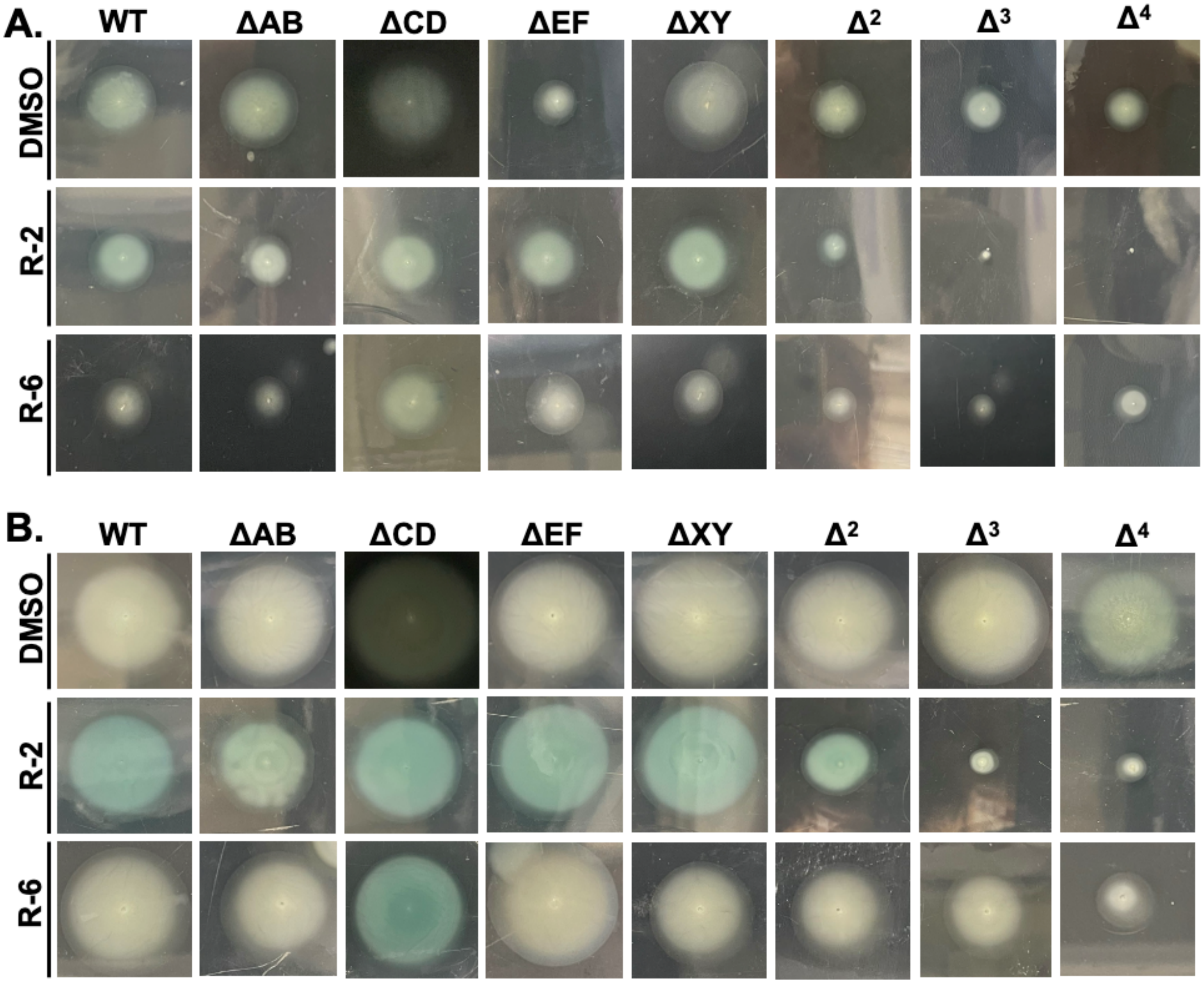


**S6 Fig: Swimming Images.** A. Representative images for 24 hr swims for DMSO, R-2 or R-6 (200 µM) treatment with each efflux pump deficient strain. B. Representative images for 48 hr swims for DMSO, R-2 or R-6 (200 µM) treatment with each efflux pump deficient strain. All experiments were completed in biological triplicate.

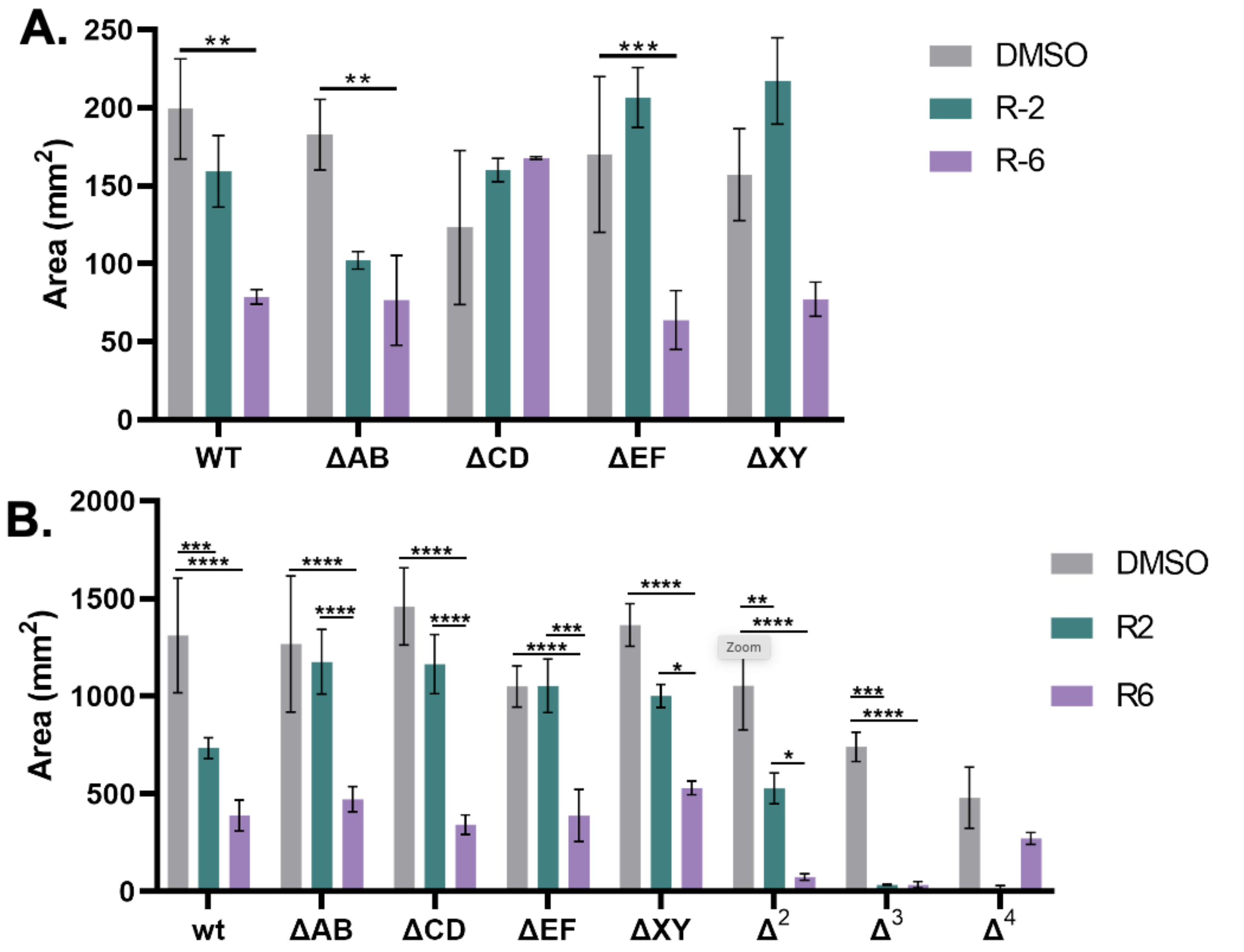


**S7 Fig: Quantified Swimming Area.** A. Calculated area for each swim at 24 hr for DMSO, R-2 or R-6 (200 µM) treated efflux pump deficient strains. Statistical significance determined by one-way ANOVA Brown-Forsythe and Welch in Prism Graphpad 9.5.0. p-value ** <0.005, ***<0.0005. B. Calculated area for each swim at 48 hr for DMSO, R-2 or R-6 (200 µM) treated efflux pump deficient strains. Statistical significance determined by two-way ANOVA in Prism Graphpad 9.5.0. p-value * <0.01, ** <0.005, ***<0.0005, ****<0.0001.

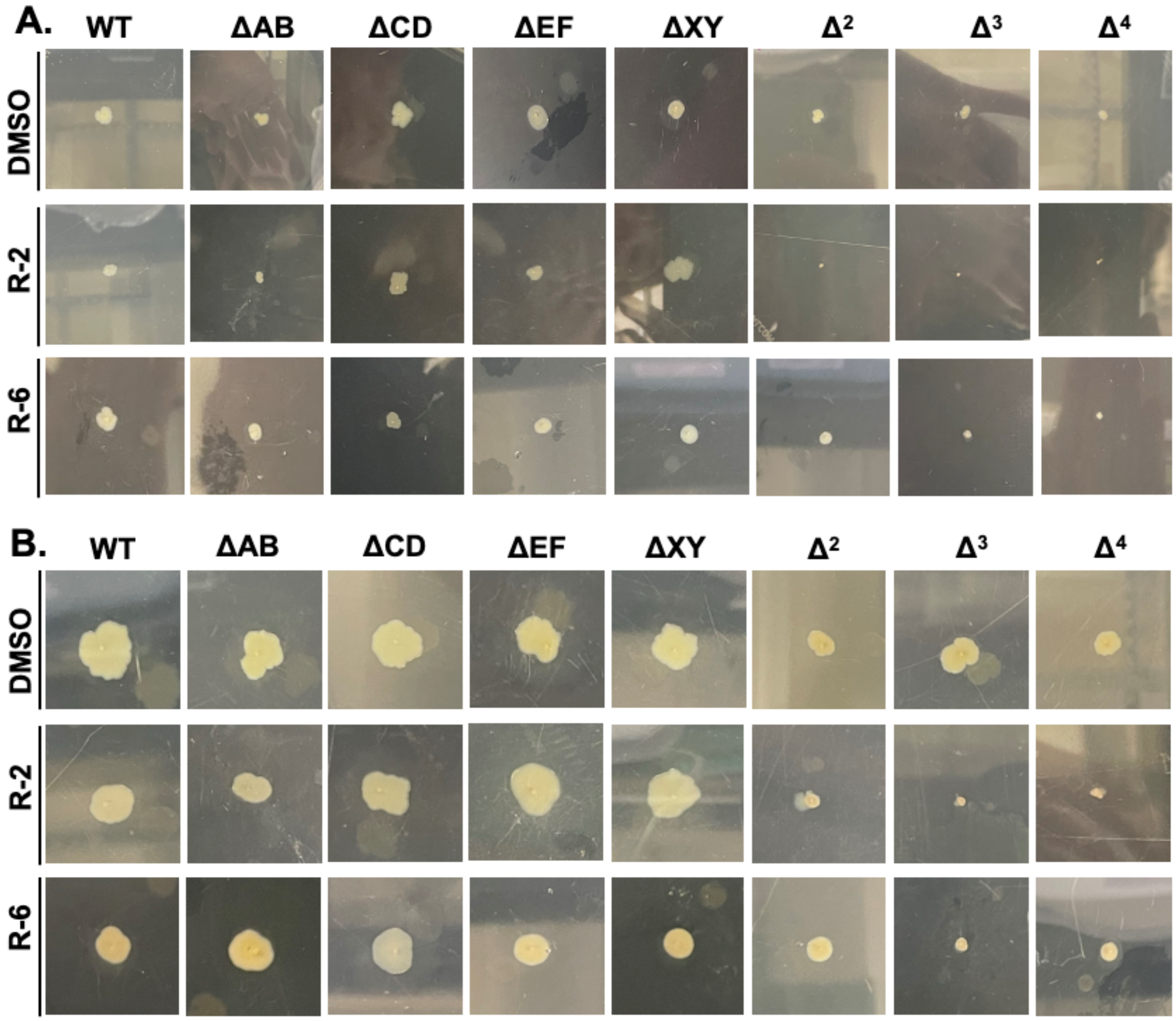


**S8 Fig: Twitching Images.** A. Representative twitching images at 24 hr for each efflux pump deficient strain of PA14 treated with DMSO, R-2 or R-6 (200 μM). B. Representative twitching images at 48 hr for each efflux pump deficient strain of PA14 treated with DMSO, R-2 or R-6 (200 μM). All experiments were completed in biological triplicate.

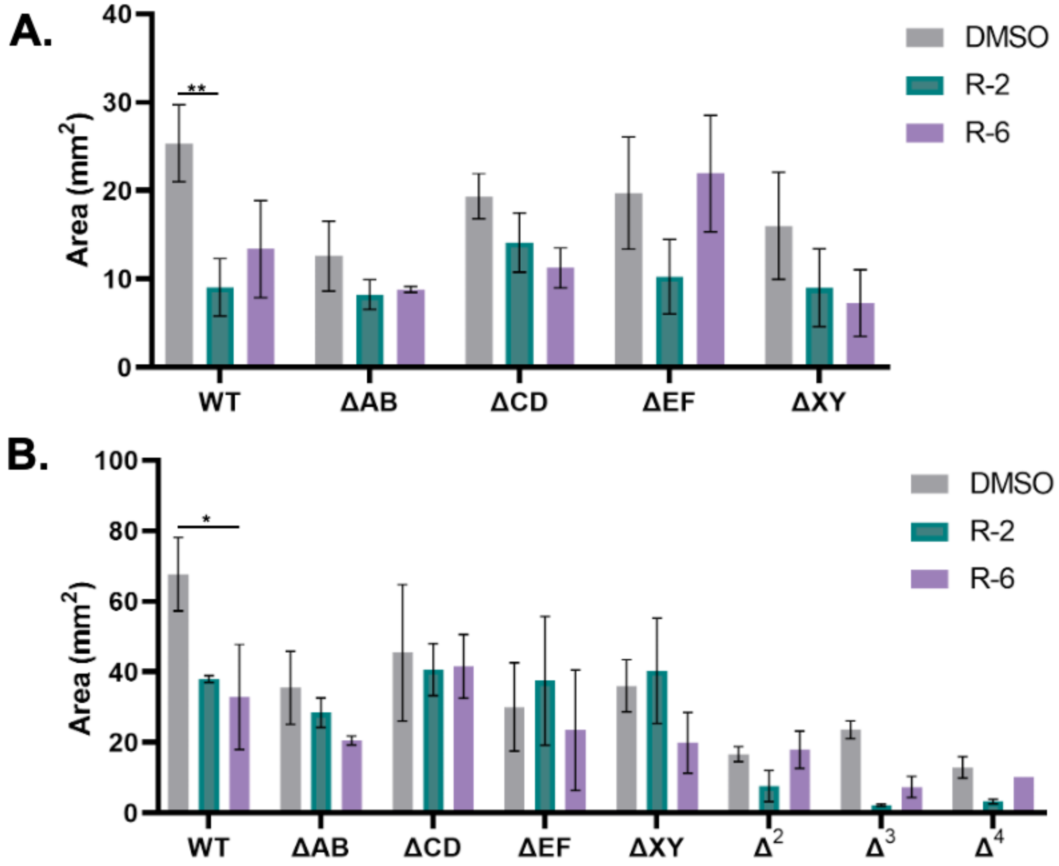


**S9 Fig: Quantified Twitching Area.** A. Representative twitch images at 48 hr for each efflux pump deficient strain of PA14 treated with DMSO, R-2 or R-6 (200 µM). B. Calculated area for each twitch at 48 hr for DMSO, R-2 or R-6 treated efflux pump deficient strains. Statistical significance determined by two-way ANOVA in Prism Graphpad 9.5.0. p-value * <0.01, ** <0.005.

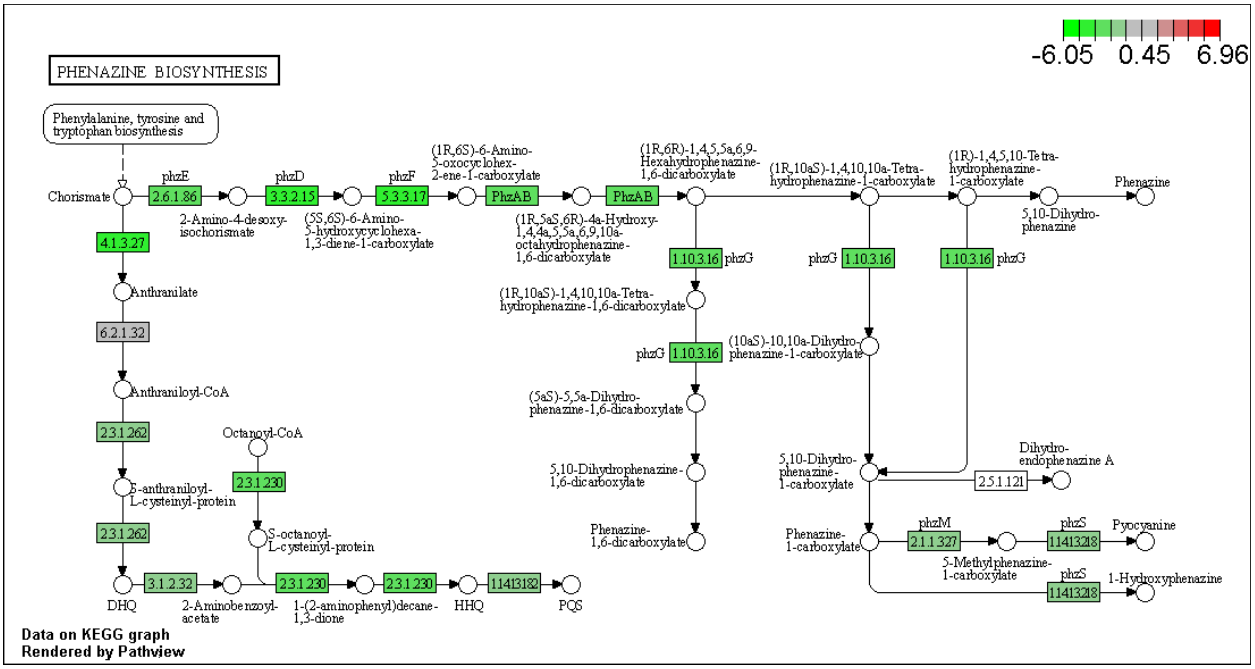


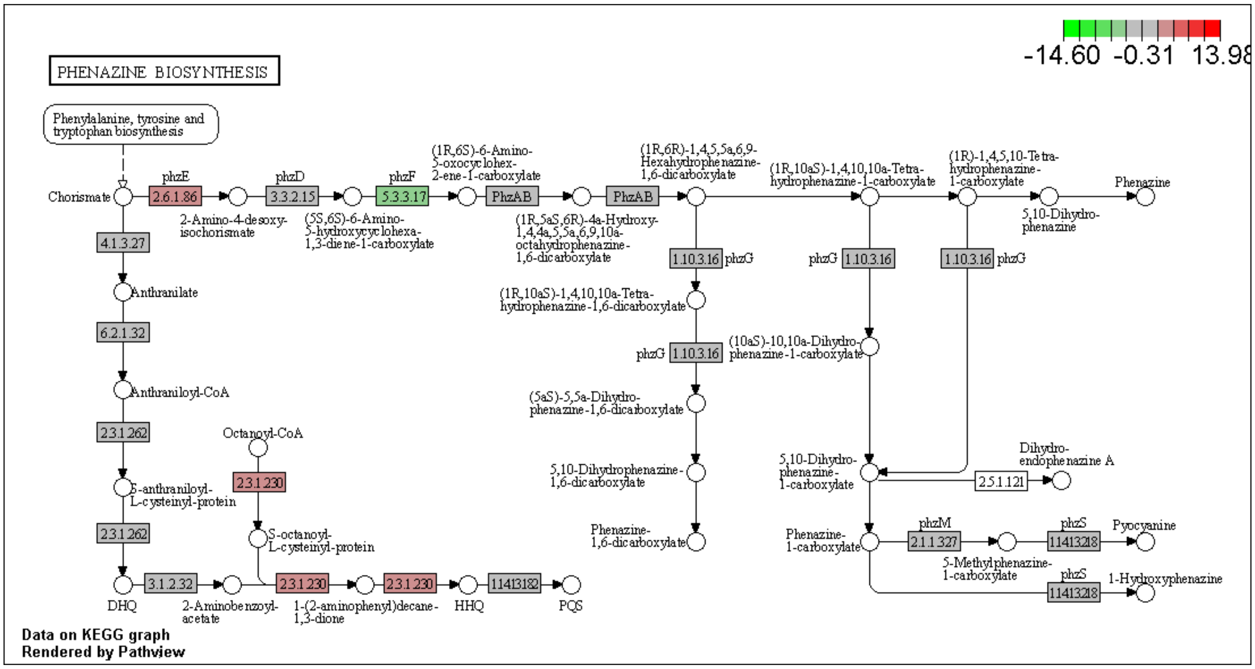


**S10 Fig: Pathview Pathway R-6 Phenazine Biosynthesis.** Canonical signaling pathway (pau00405 Phenazine Biosynthesis) with gene data from R-6E and R-6L, top and bottom dataset respectively, compared to DMSO control.

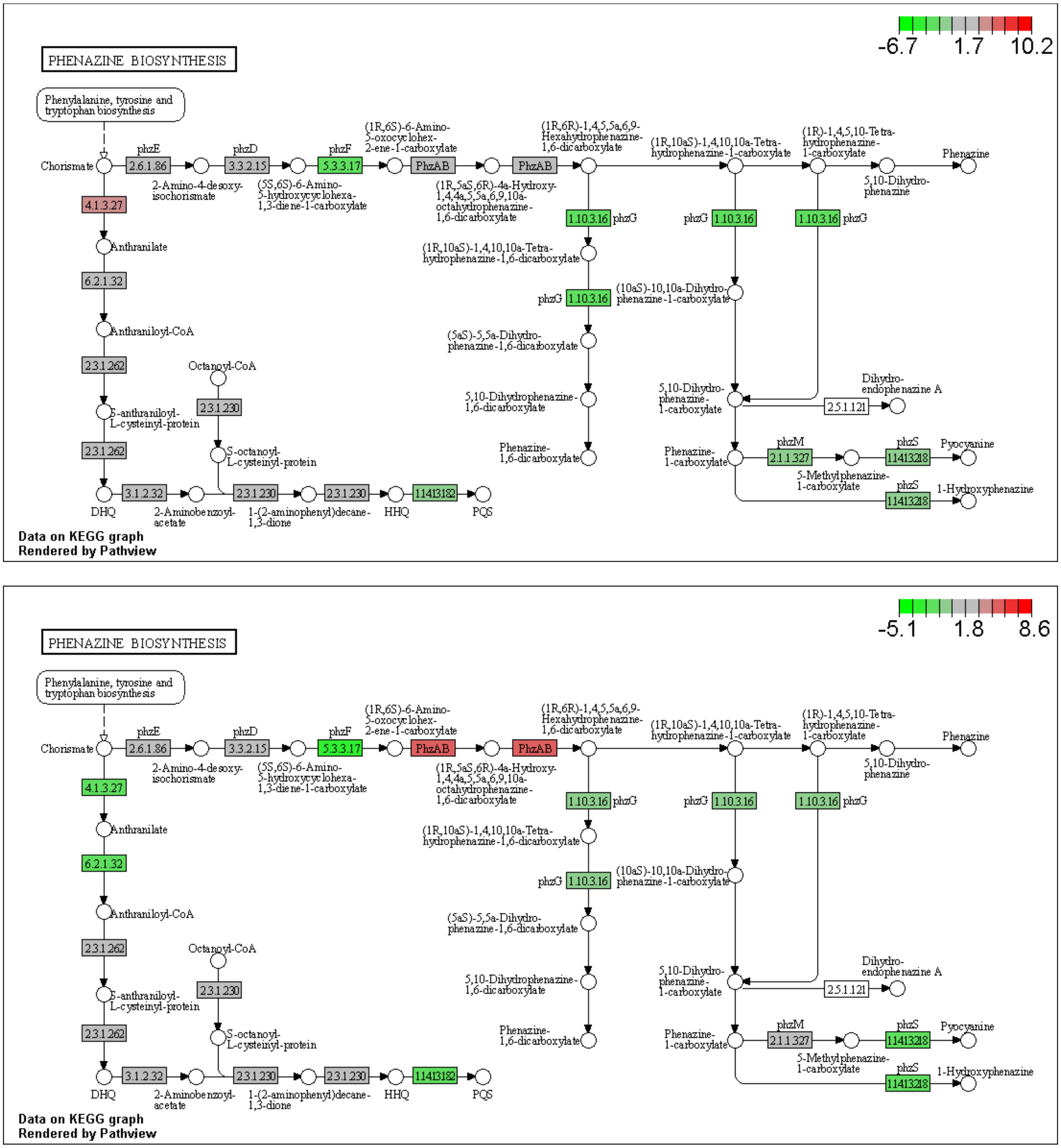


**S11 Fig: Pathview Pathway R-2 Phenazine Biosynthesis.** Canonical signaling pathway (pau00405 Phenazine Biosynthesis) with gene data from R-2E and R-2L, top and bottom dataset respectively, compared to DMSO control.

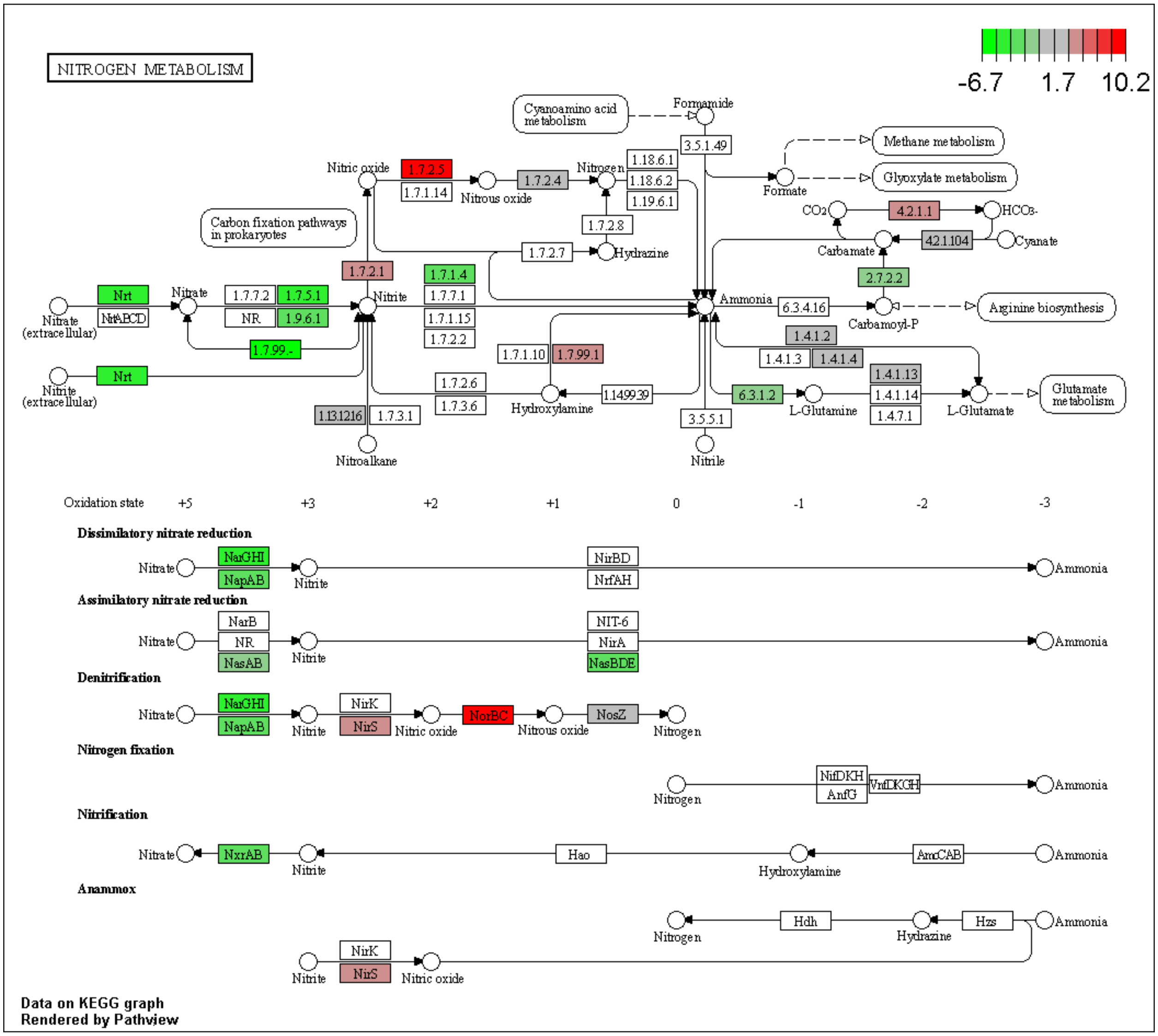


**S12 Fig: Pathview Pathway R-2E Nitrate Respiration.** Canonical signaling pathway (pau00910 Nitrogen Metabolism) with gene data from R-2E dataset compared to DMSO control.

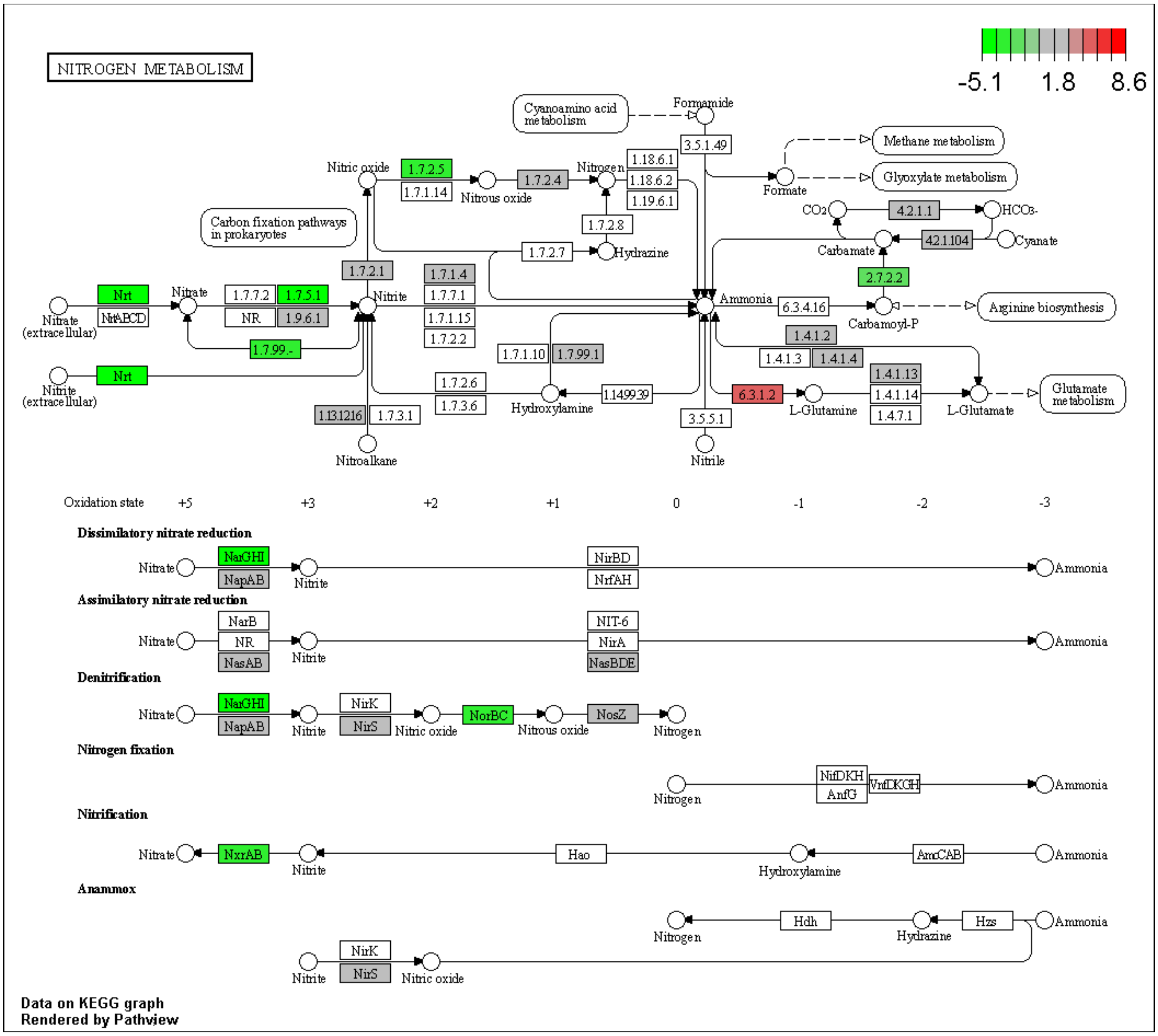


**S13 Fig: Pathview Pathway R-2L Nitrate Respiration.** Canonical signaling pathway (pau00910 Nitrogen Metabolism) with gene data from R-2L dataset compared to DMSO control.

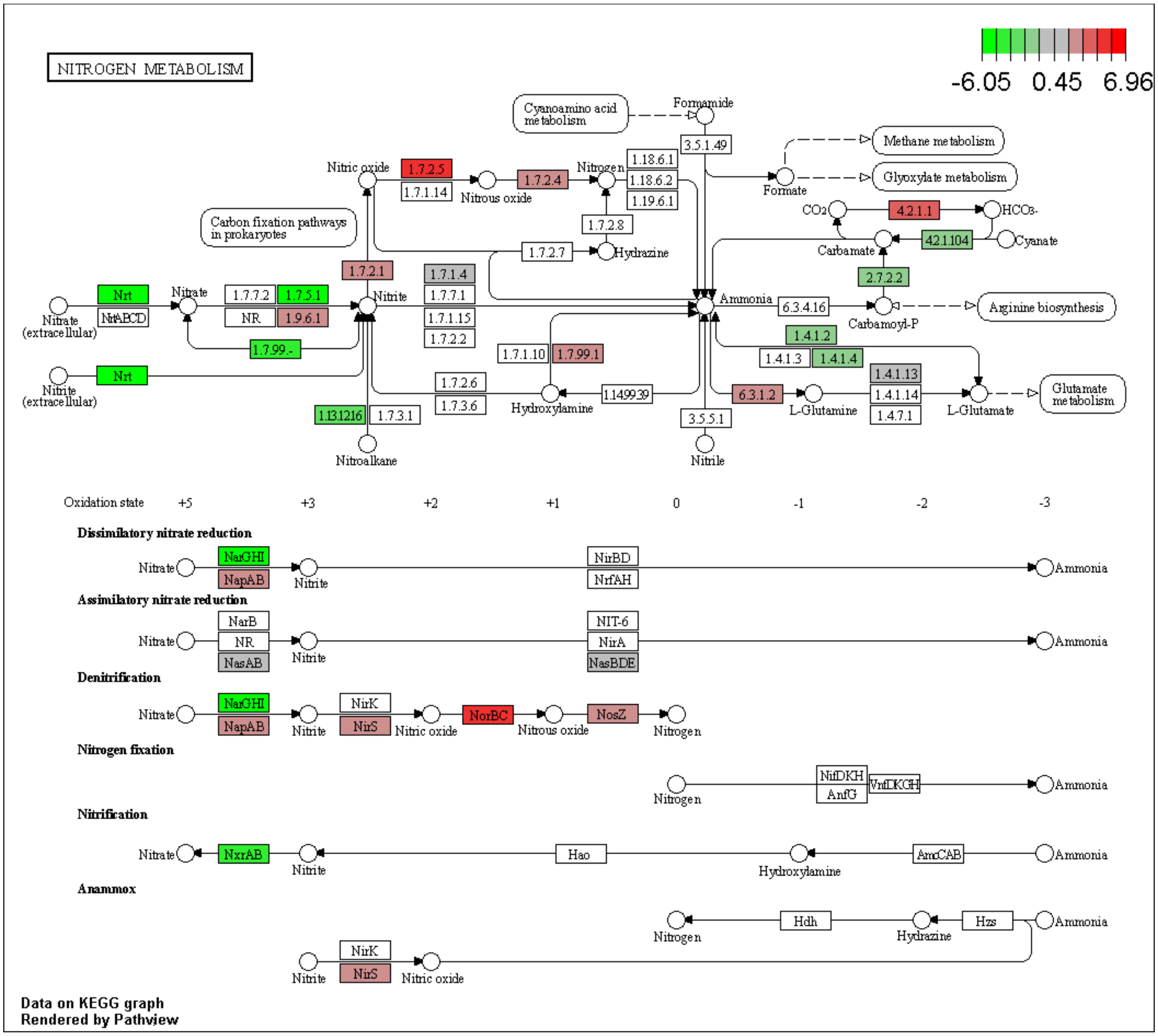


**S14 Fig: Pathview Pathway R-6E Nitrate Respiration.** Canonical signaling pathway (pau00910 Nitrogen Metabolism) with gene data from R-6E dataset compared to DMSO control.

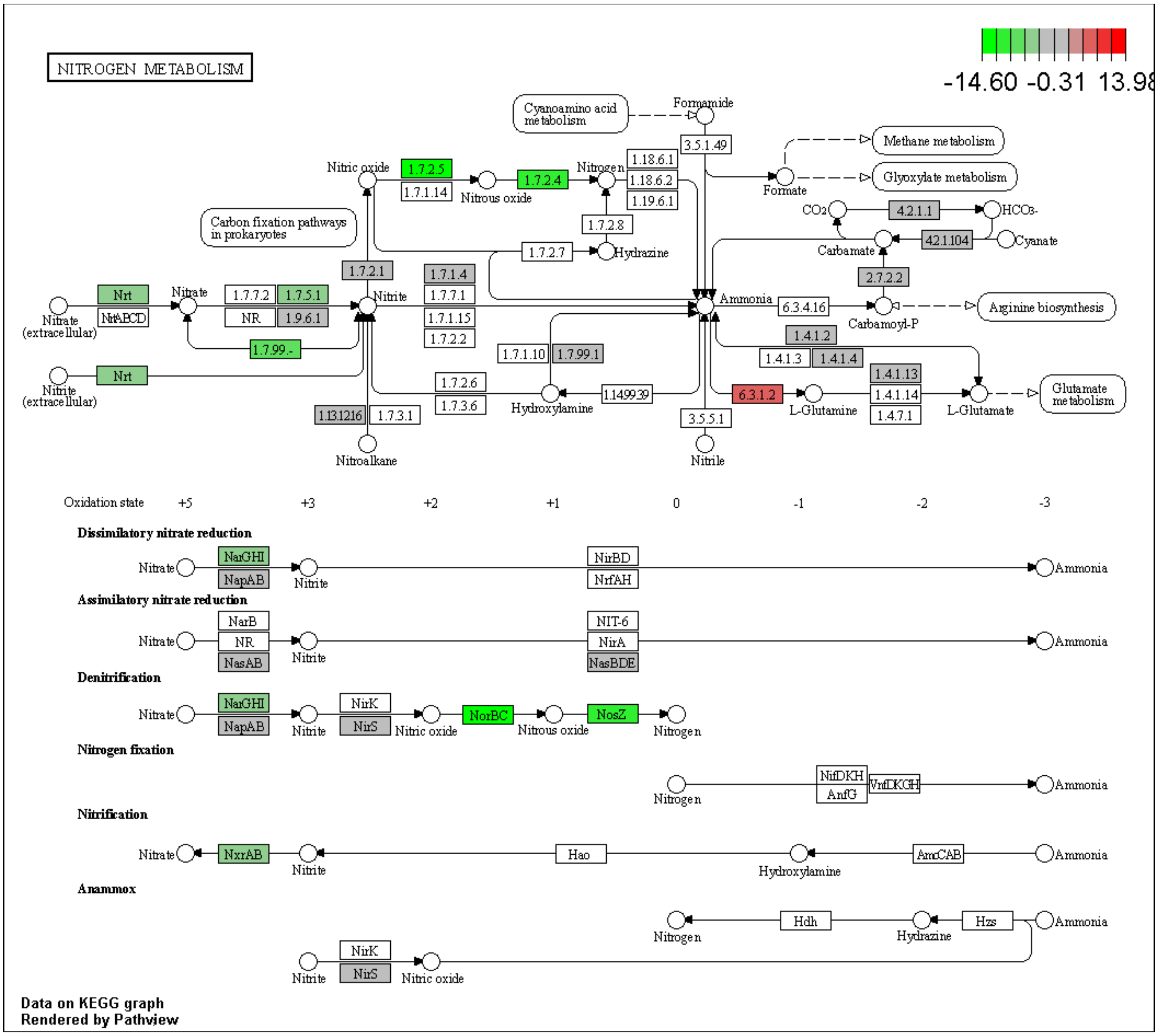


**S15 Fig: Pathview Pathway R-6L Nitrate Respiration.** Canonical signaling pathway (pau00910 Nitrogen Metabolism) with gene data from R-6L dataset compared to DMSO control.

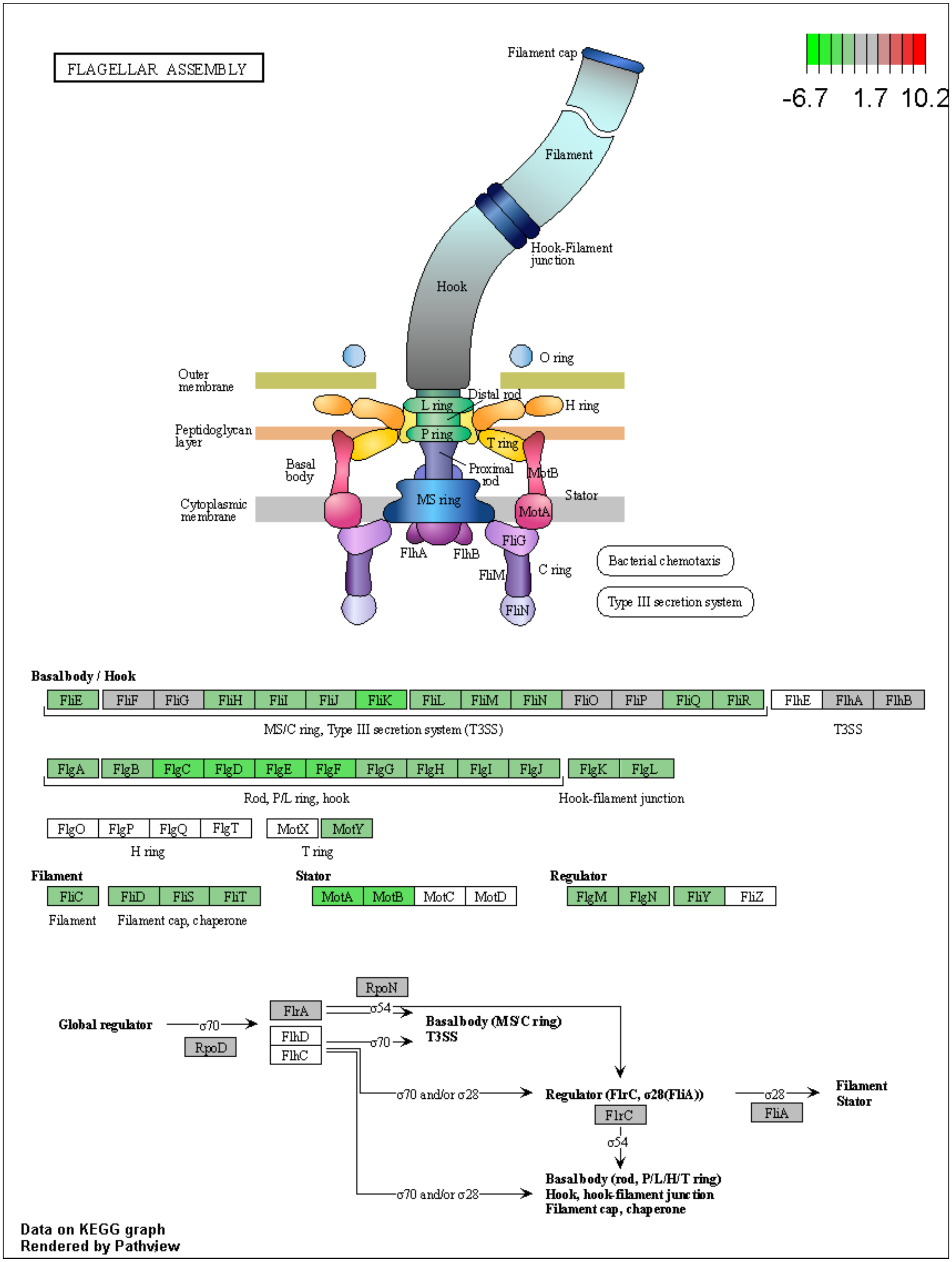


**S16 Fig: Pathview Pathway R-2E Flagellar Assembly.** Canonical signaling pathway (pau02040 Flagellar Assembly) with gene data from R-2E dataset compared to DMSO control.

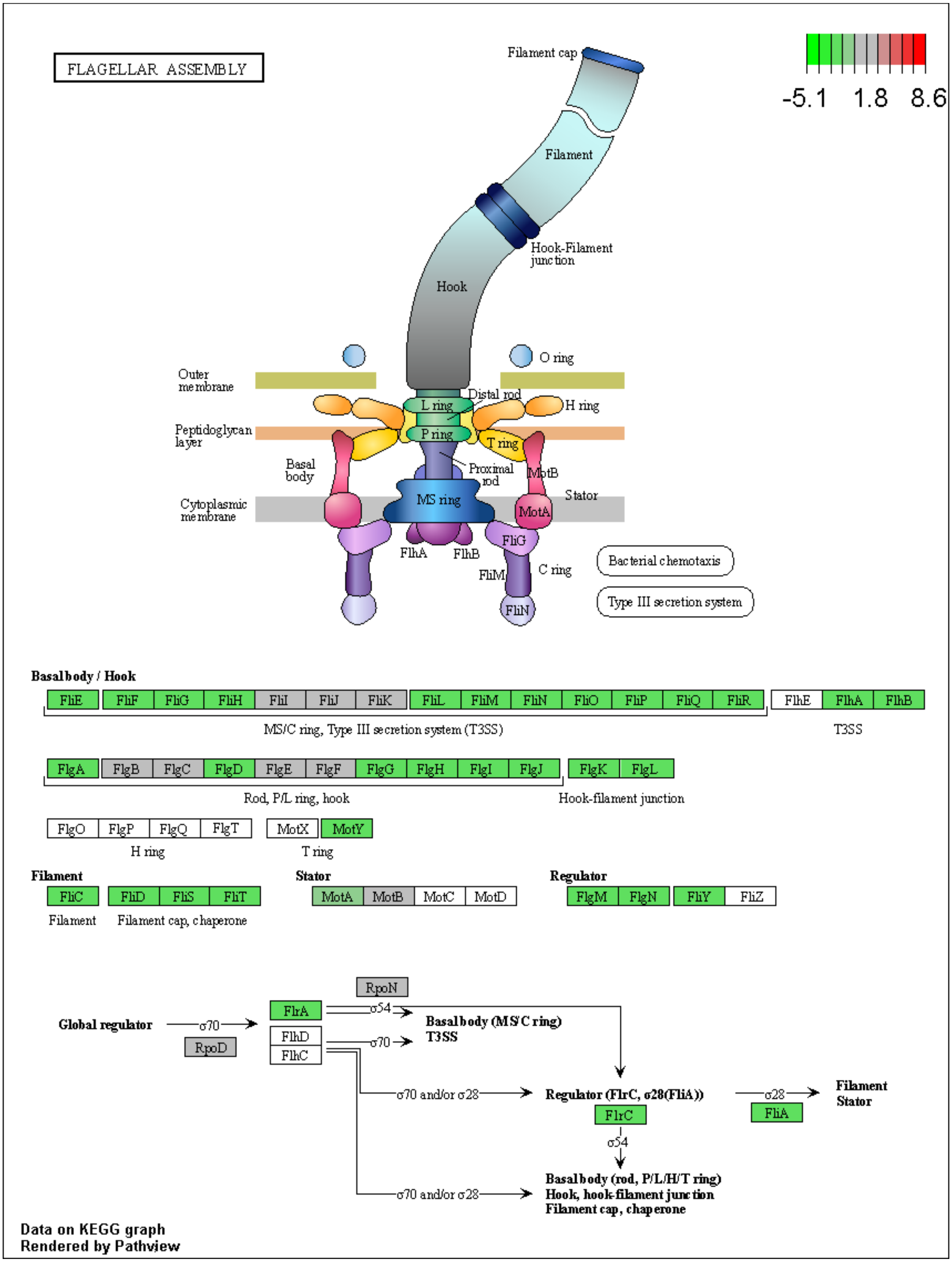


**S17 Fig: Pathview Pathway R-2L Flagellar Assembly.** Canonical signaling pathway (pau02040 Flagellar Assembly) with gene data from R-2L dataset compared to DMSO control.

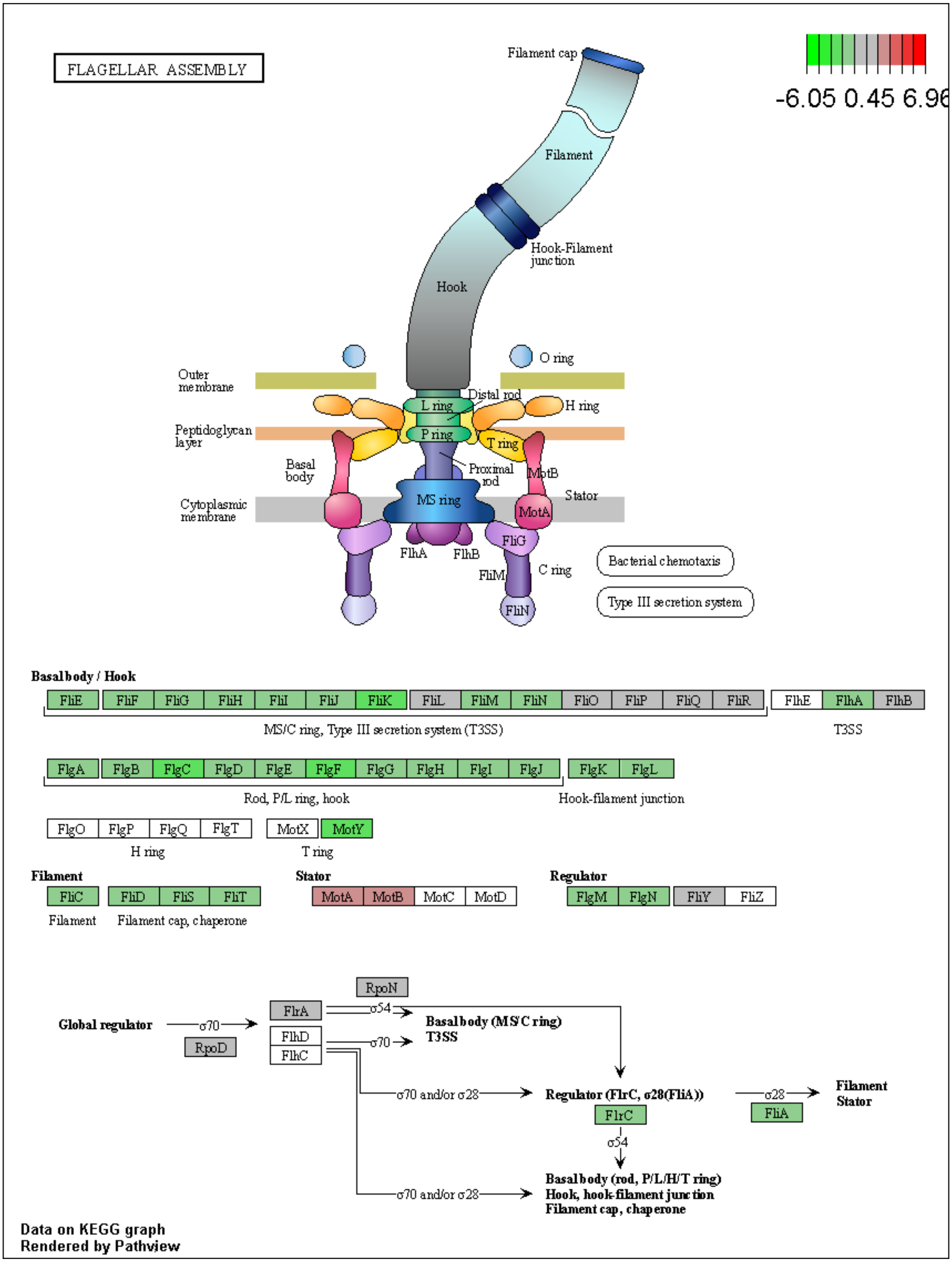


**S18 Fig: Pathview Pathway R-6E Flagellar Assembly.** Canonical signaling pathway (pau02040 Flagellar Assembly) with gene data from R-6E dataset compared to DMSO control.

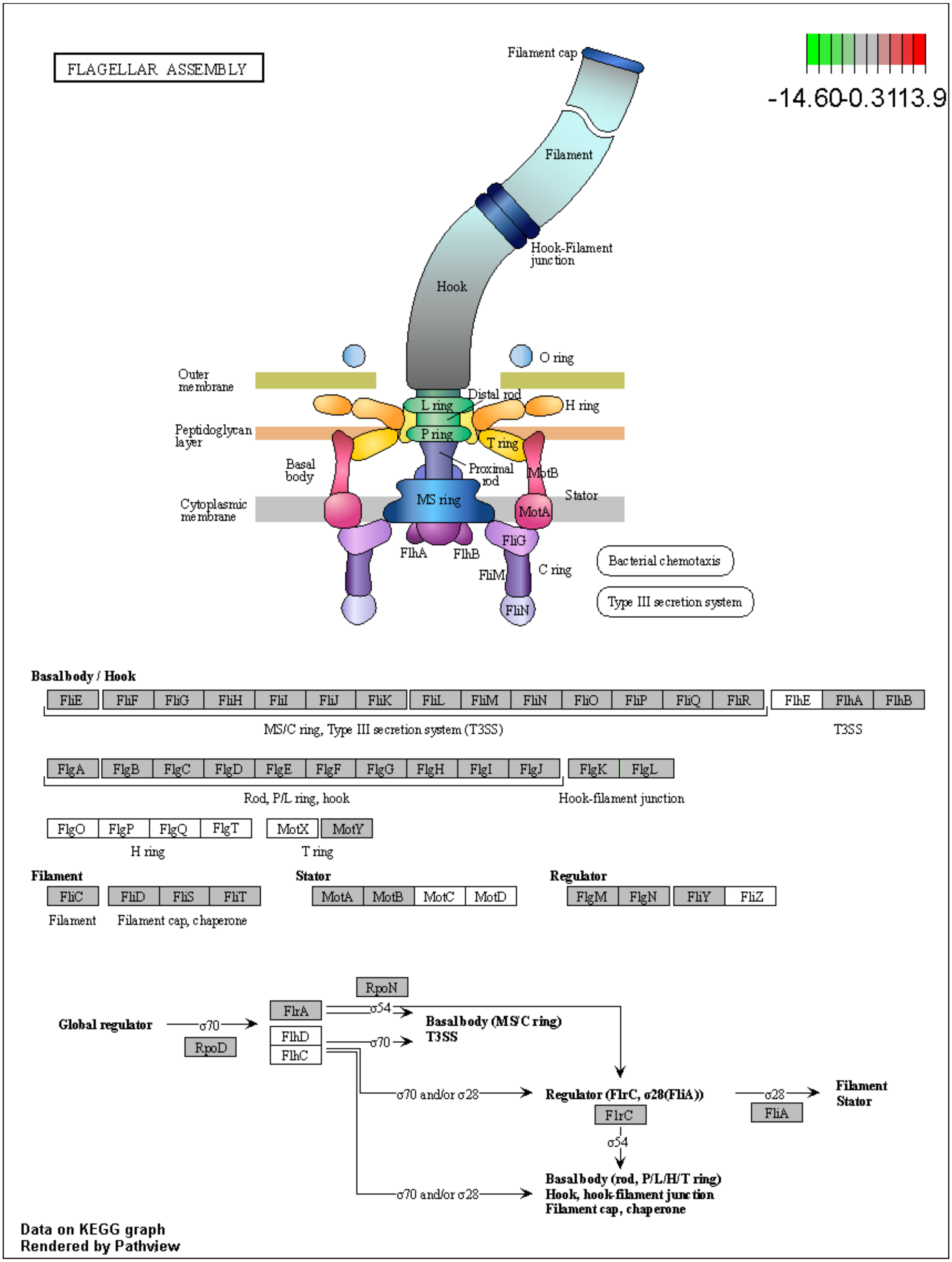


**S19 Fig: Pathview Pathway R-6L Flagellar Assembly.** Canonical signaling pathway (pau02040 Flagellar Assembly) with gene data from R-6L dataset compared to DMSO control.

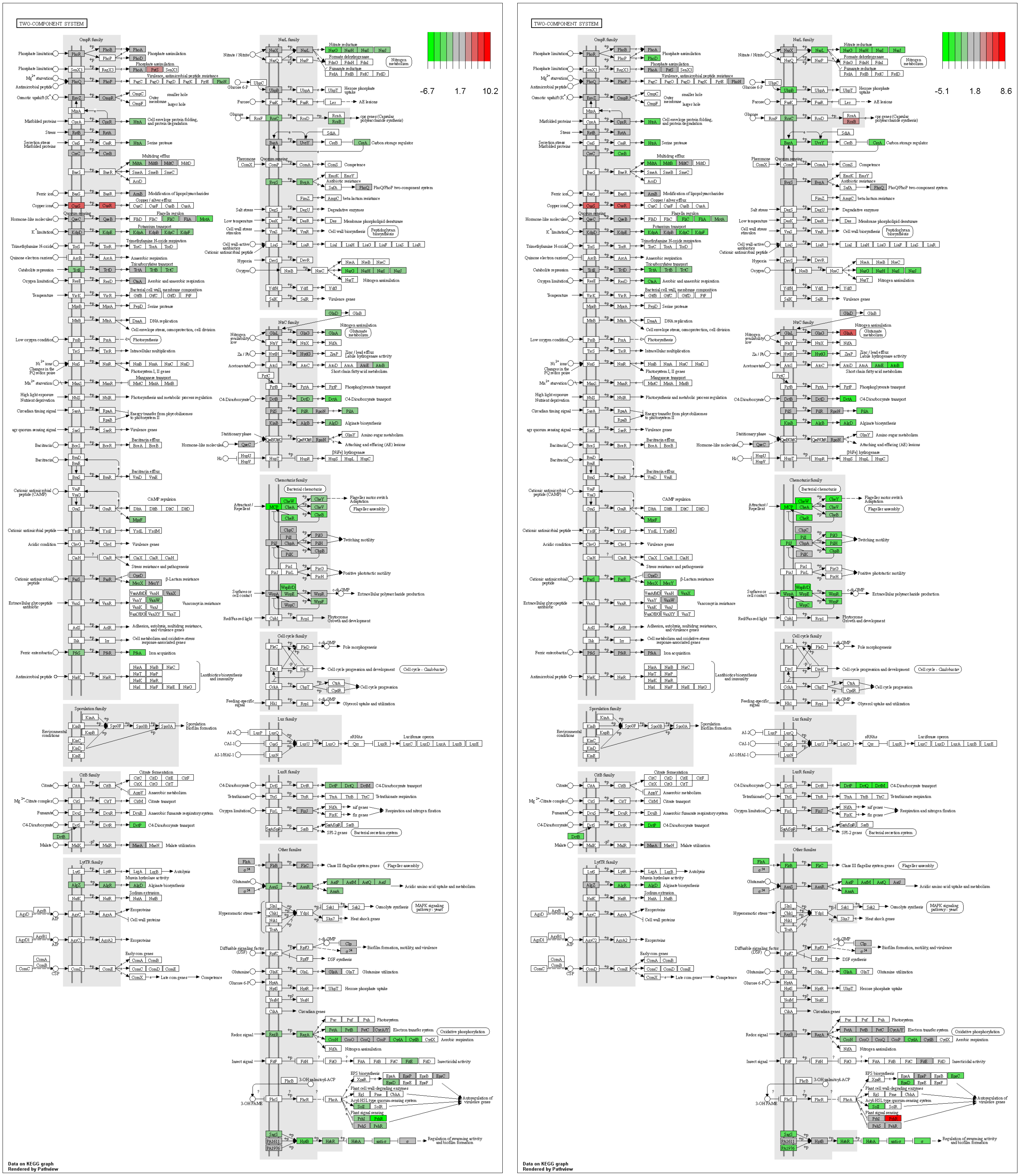


**S20 Fig: Pathview Pathway R-2 Two-Component Systems.** Canonical signaling pathway (pau02020 Two Component Systems) with gene data from R-2E and R-2L datasets, left and right respectively, compared to DMSO control.

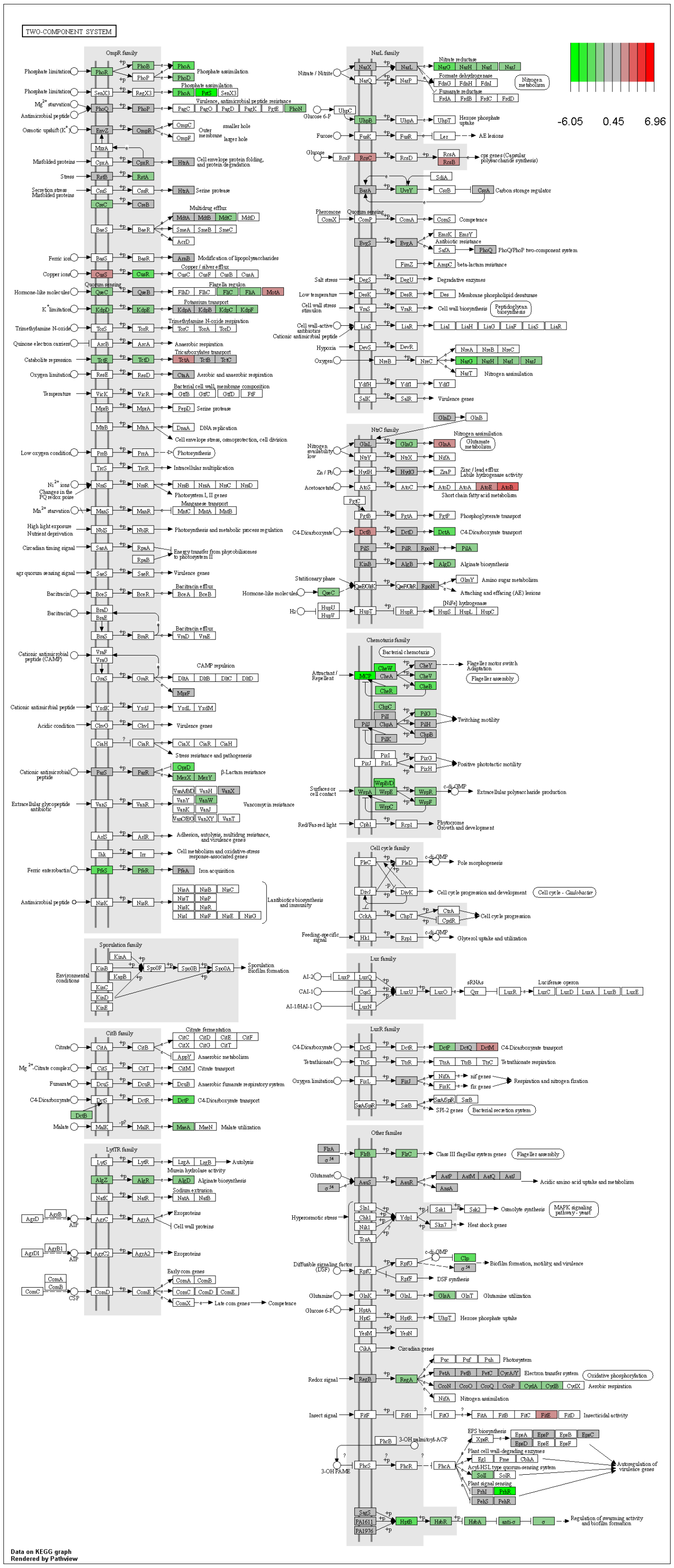


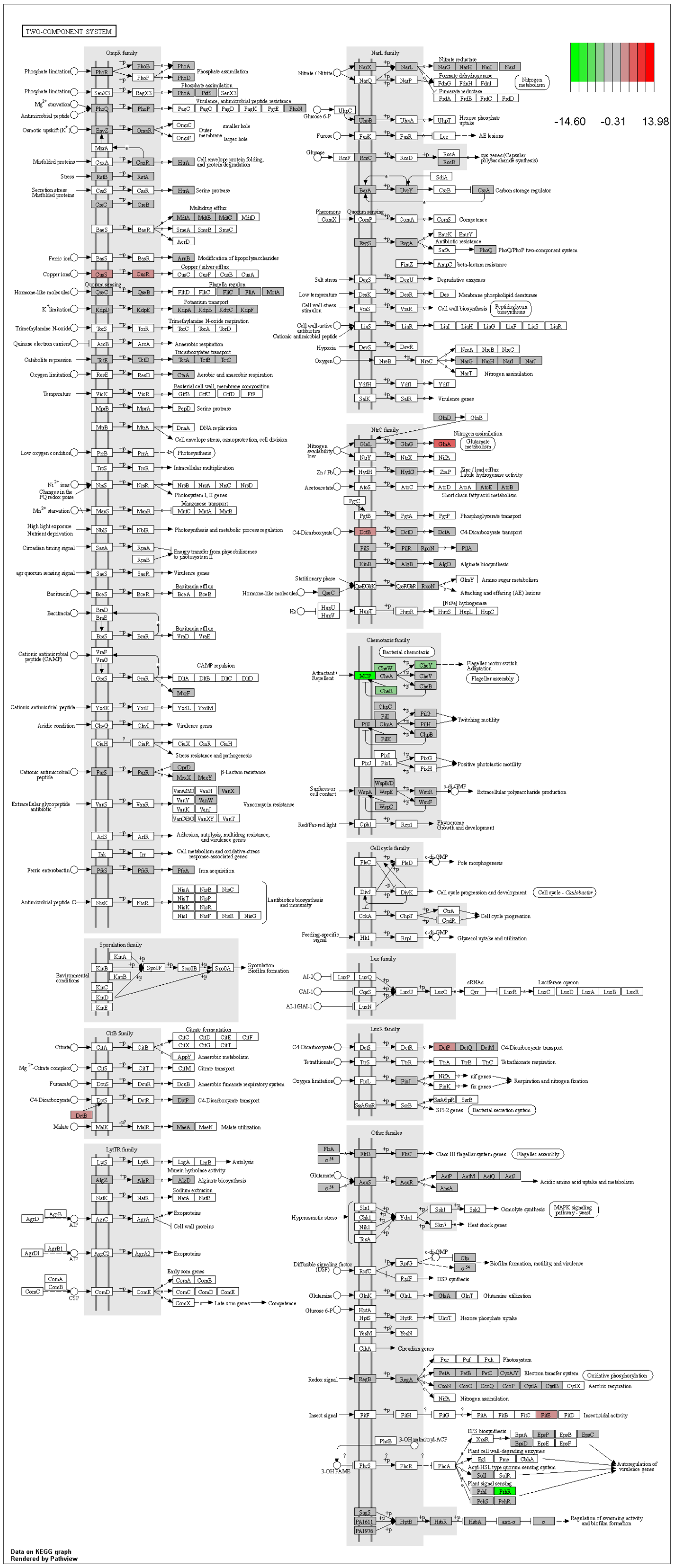


**S21 Fig: Pathview Pathway R-6 Two-Component Systems.** Canonical signaling pathway (pau02020 Two Component Systems) with gene data from R-6E and R-6L datasets, left and right respectively, compared to DMSO control.

**References**

1. Gibson DG, Young L, Chuang RY, Venter JC, Hutchison CA, 3rd, Smith HO. Enzymatic assembly of DNA molecules up to several hundred kilobases. Nat Methods. 2009;6(5):343-5.
2. The Inoue Method for Preparation and Transformation of Competent *E. coli* "Ultra-Competent" Cells: Cold Spring Harbor Laboratory Press; 2006.
3. Miller JH. Experiments in Molecular Genetics: Cold Spring Harbor Laboratory; 1972.
